# Cephalopod Genome Expansion Drives Broader Reflectin Domain Boundaries

**DOI:** 10.64898/2026.04.14.718489

**Authors:** Olivia J. Armendarez, Duncan Q. Bower, Kaitlyn R. Flynn, Michael R. Bergman, Caroline B. Albertin, Leila F. Deravi

## Abstract

As more genome libraries become accessible across multiple cephalopod species, long standing assumptions about the molecular basis of their dynamic optical systems are being revisited. For example, cephalopod-specific reflectin proteins now appear far more variable as newly annotated sequences diverge from the canonical features that have long defined this protein family. In this perspective, we discuss these nuances and introduce a theoretical framework that expands the reflectin domain classification while preserving specificity across 141 known and previously uncharacterized reflectin sequences from ten cephalopod species. By combining this broadened domain architecture with quantitative, bioinformatics-driven metrics, we establish a generalizable framework that reliably identifies reflectins across their full sequence diversity, allowing for deeper structural comparisons and future functional discovery.

## INTRODUCTION

Reflectins are self-assembling proteins found in cephalopod mollusks that have attracted substantial scientific interest over the past two decades due to their capacity to generate structure-dependent iridescent optical properties.^1-12^ Their hierarchical assembly and resultant photonic properties have been the subject of extensive investigation across multiple disciplines including biomaterials^13,14^ used to produce adaptive coatings,^15,16^ fibers for color-changing fabrics,^17^ transducing elements,^18,19^ infrared-concealment,^20,21^ and more recently modified cell properties.^22,23^ Recently, the emergence of genomic data from multiple species of cephalopods has uncovered dozens of newly discovered reflectin subtypes and isoforms, necessitating an investigation into whether these differences influence unique optical functions that may similarly be harnessed for materials applications.

Unlike other organisms that exhibit tunable iridescence,^24,25^ cephalopods have evolved reflectin-based structural colorants present throughout their dermal tissue.^26-29^ These proteins are observed within the chromatophores, localized within the sheath cells of these dynamic organs.^30-32^ They are also found in leucophores, assembled as nanostructures^11,33-36^ that produce diffuse white patches. A notable exception is a rare case in female *D. opalescens* which transitions between transparent and bright white.^8^ In the iridosomes, reflectin assembles as lamellae (100-200 nm in diameter) that reflect narrowband light. Interestingly, reflectins within some iridophores can be static or dynamic.^7,10,12,37-39^ Loliginids (belonging to the family Loliginidae) are squid species known to have dynamic iridophores that exhibit reversible changes in reflected color.^40^ These responsive iridophores can be localized to regions of the skin that can be stimulated with exogenous acetylcholine, as demonstrated in *D. opalescens*.^38^ However, the connection between reflectin variants and the dynamic nature of the iridophores they occupy remains unclear. It is also unknown why some cephalopods are endowed with dynamic iridophores while others are not.

Recently, photophores in nocturnal or deep-sea cephalopod species have also gained interest for their ability to generate light, often in flashes.^41^ Bacterial photophores, observed in cephalopods, utilize a symbiotic relationship with microbes growing in specialized chambers within their host.^42-44^ In these organs, reflectins organized as flat platelets are also used to reflect and direct the light produced.^41,44^ Some photophores even have opaque structures surrounding them that can be shuttered to emit only flashes of light.^5^ Based on these observations, it is clear that reflectins offer a diversity in structures and functions that are essential to cephalopods in unique ways, yet it remains unclear why a single protein class has evolved to produce so many specific functions across multiple organs within these animals. In this work, we consider the structure and composition of reported and uncharacterized reflectins from squid (longfin inshore squid, *Doryteuthis pealeii*; opalescent inshore squid, *Doryteuthis opalescens*; Hawaiian bobtail squid, *Euprymna scolopes*; Atlantic giant squid, *Architeuthis dux*; hummingbird bobtail squid, *Euprymna berryi*), cuttlefish (common cuttlefish, *Sepia officinalis*; pharaoh cuttlefish, *Sepia pharaonis*), and octopus (greater argonaut octopus, *Argonauta argo*; California two-spot octopus, *Octopus bimaculoides*; common octopus, *Octopus vulgaris*) and use this information to extrapolate correlations between domain-linker organization and potential function.

### REFLECTIN SEQUENCE VARIATION AMONG CEPHALOPODS

To explore potential correlations between sequence and function, we gathered 141 known and uncharacterized reflectin sequences from ten cephalopod species available through both UniProt and the Marine Biological Laboratory (**Table 1, Table S1)**. Of these, 32 sequences belong to *E. scolopes*, 21 to *D. pealeii*, 4 to *D. opalescens*, 14 to *A. dux*, 21 to *E. berryi*, 9 to *S. pharaonis*, 17 to *S. officinalis*, 6 to *A. argo*, 8 to *O. bimaculoides*, and 9 to *O. vulgaris*. Initial alignments of reflectin sequences within each species using Clustal Omega^45,46^ revealed varying conservation (0.7%-99%) across all species (**Table S2**). To explore this variability further, we compared sequence similarities within *E. scolopes, D. pealeii*, and *O. bimaculoides* (**Figure 1**). The repeating pattern in *E. scolopes* differs from that observed in *D. pealeii*. Because reflectins belonging to the *Octopus* genus are not currently well-classified, we first identified putative reflectin sequences manually using a highly conserved reflectin sequence from the pharaoh cuttlefish *S. pharaonis* (MMEPMSRMTMDFQGRYMDSQGRMV) as a query in a BLASTp search. The results included reflectin sequences from *S. officinalis* and *S. pharaonis* and two octopus species *O. vulgaris* and *O. bimaculoides* available in the Uniprot database. The sequences for *O. vulgaris* range in identity from 16.1% and 97.0% and sequences for *O. bimaculoides* range from 21.3% to 98.8% identity. These sequences also present similarities to reflectin-like domains (**Figure 1**). One section that shows the most similarity to *D. pealeii* has the consensus motif of [MF(X)_7_MD(X)_5_MD(X)_5_].

**Table 1.**
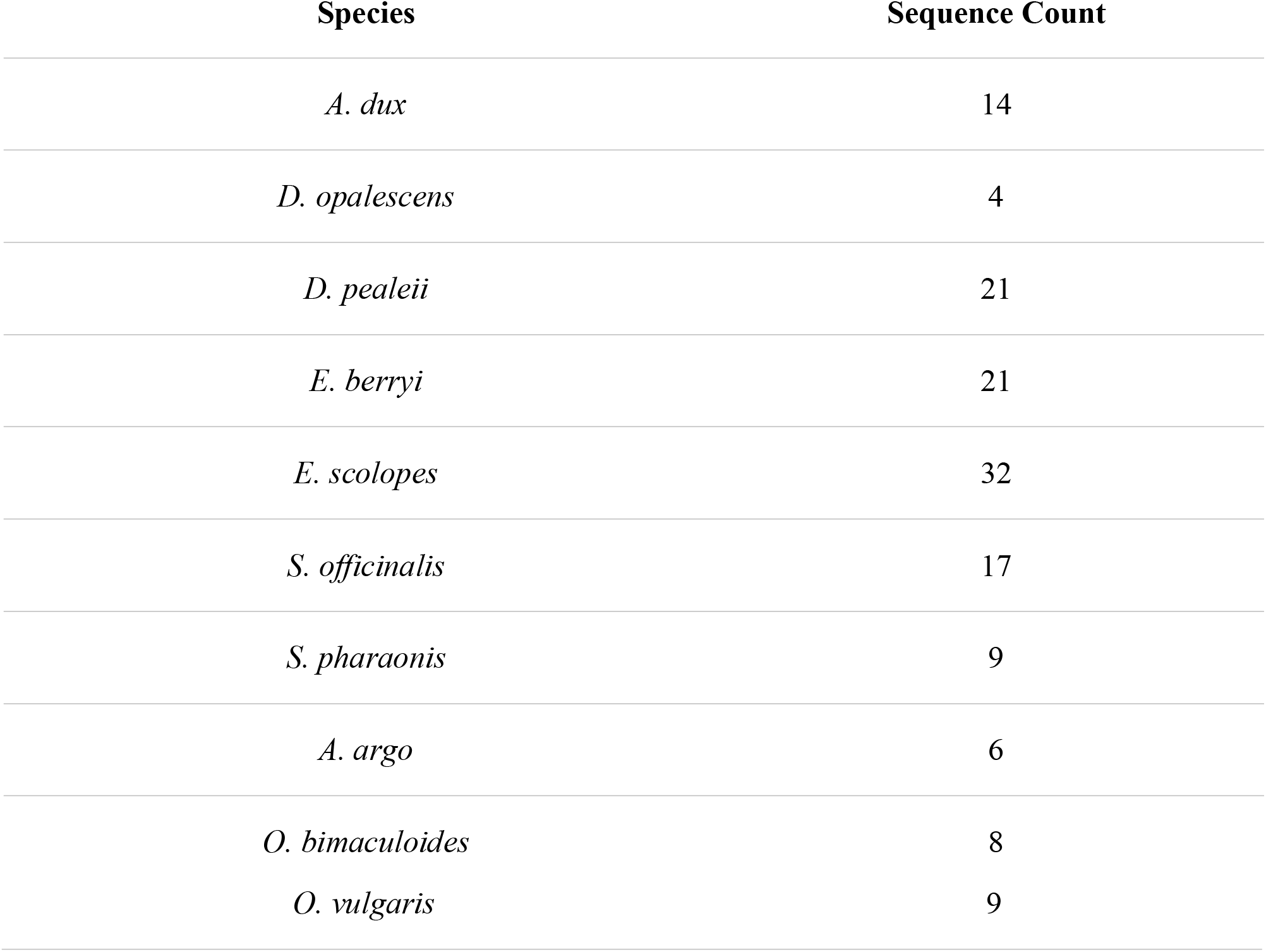
Cephalopod species with corresponding sequence count analyzed in this study.

**Figure 1.**
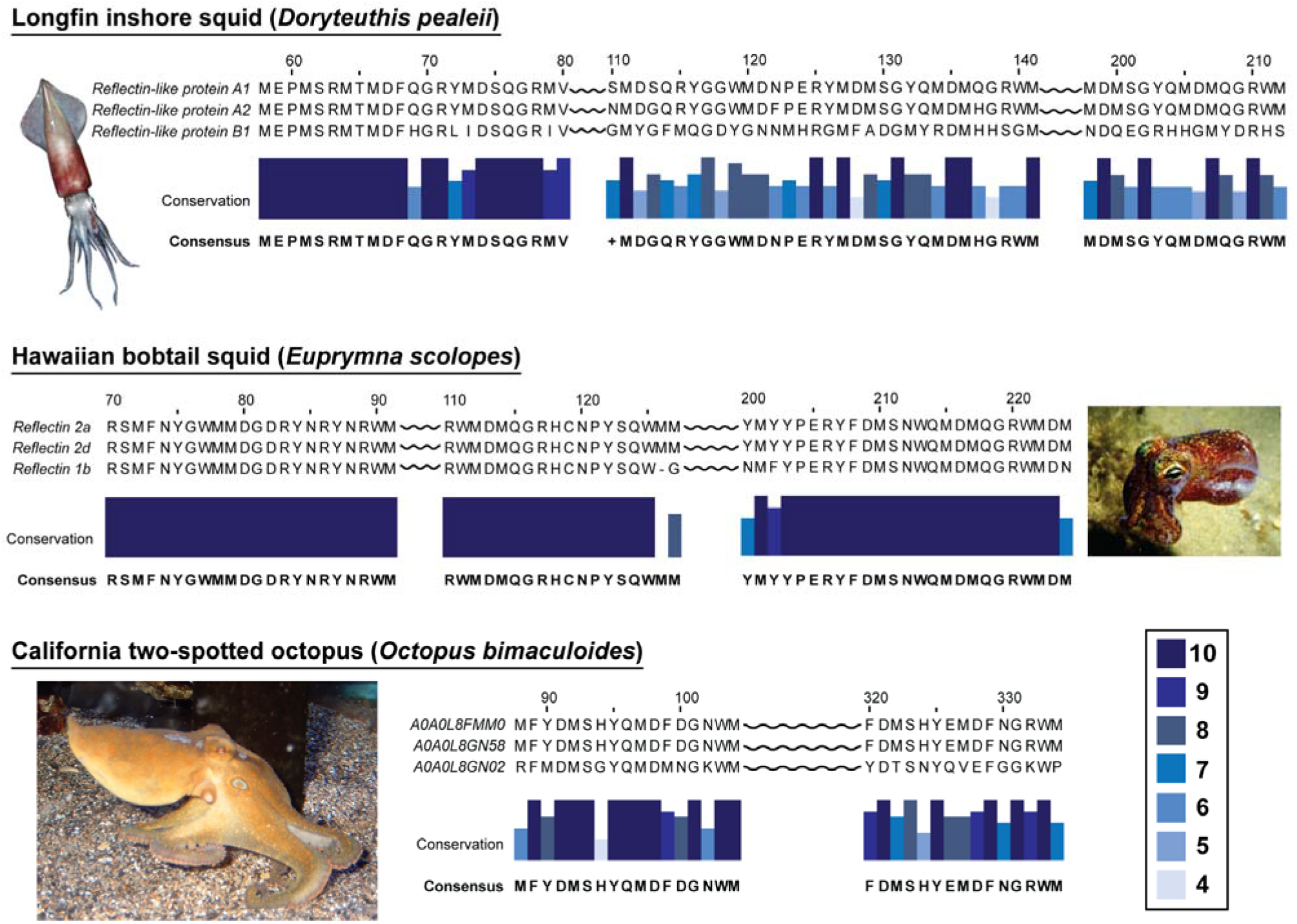
Alignment of reflectin and reflectin-like protein sequences from three cephalopods. Sequences were retrieved from the UniProt database with each being labeled according to the protein name except for *O. bimaculoides* which consisted of “uncharacterized proteins” and thus the accession numbers are shown instead. Sequences for *D. pealeii, E. scolopes*, and *O. bimaculoides* were aligned using the MUSCLE^47,48^ algorithm in Jalview.^49,50^ For *E. scolopes* and *O. bimaculoides*, the three sequences above represent the two most identical sequences, and the most dissimilar sequence, to highlight sequence diversity within a singular organism. Conservation was calculated^51^ in Jalview and illustrated here by both the shade and height of the blue bars (see legend for numerical assignments). Both conservation and the consensus sequence were determined from only the three sequences shown for each organism where each residue in 1-letter code represents the most conserved amino acid across the three sequences, and + is used when no consensus was reached. All three images were obtained from the public domain.

### RELAXING THE REFLECTIN DOMAIN DEFINITION

Based on our initial analysis, it is clear that the canonical tri-repeating motif originally proposed from the *E. scolopes* analysis^7^ might not be inclusive for the reflectins identified across all species. In the original study of seven sequences found in *E. scolopes*, reflectin domains followed the repeat pattern [M/FD(X)_5_MD(X)_5_MD(X)_3/4_], spaced with cationic linkers.^7,52^ However, this domain structure showed poor coverage across all sequences in our expanded dataset (**Figure 2A**). To expand this definition, we used the number of repeating methionine motifs and a semi-restricted range of residues in the “Z” position immediately following each methionine to establish criteria for categorizing a “reflectin domain”. Specifically, we defined a reflectin domain as a pattern of methionine (“M”)-non-methionine (“Z”) repeating at least three times, where Z is aspartic acid (D), serine (S), glutamine (Q), tyrosine (Y), asparagine (N), phenylalanine (F), or histidine (H). Between methionine repeats, a spacer (“X”) of up to five residues was permitted to maintain consistency with the canonical reflectin definition. Although we observed spacers longer than five residues in some sequences, we limited our analysis to X≤5 for this reason. The N-terminal initiator methionine was excluded from this analysis. For each domain, we included five residues downstream of the final methionine to capture the terminal (X)_3/4_ spacer characteristic of *E. scolopes* reflectin domains.

**Figure 2.**
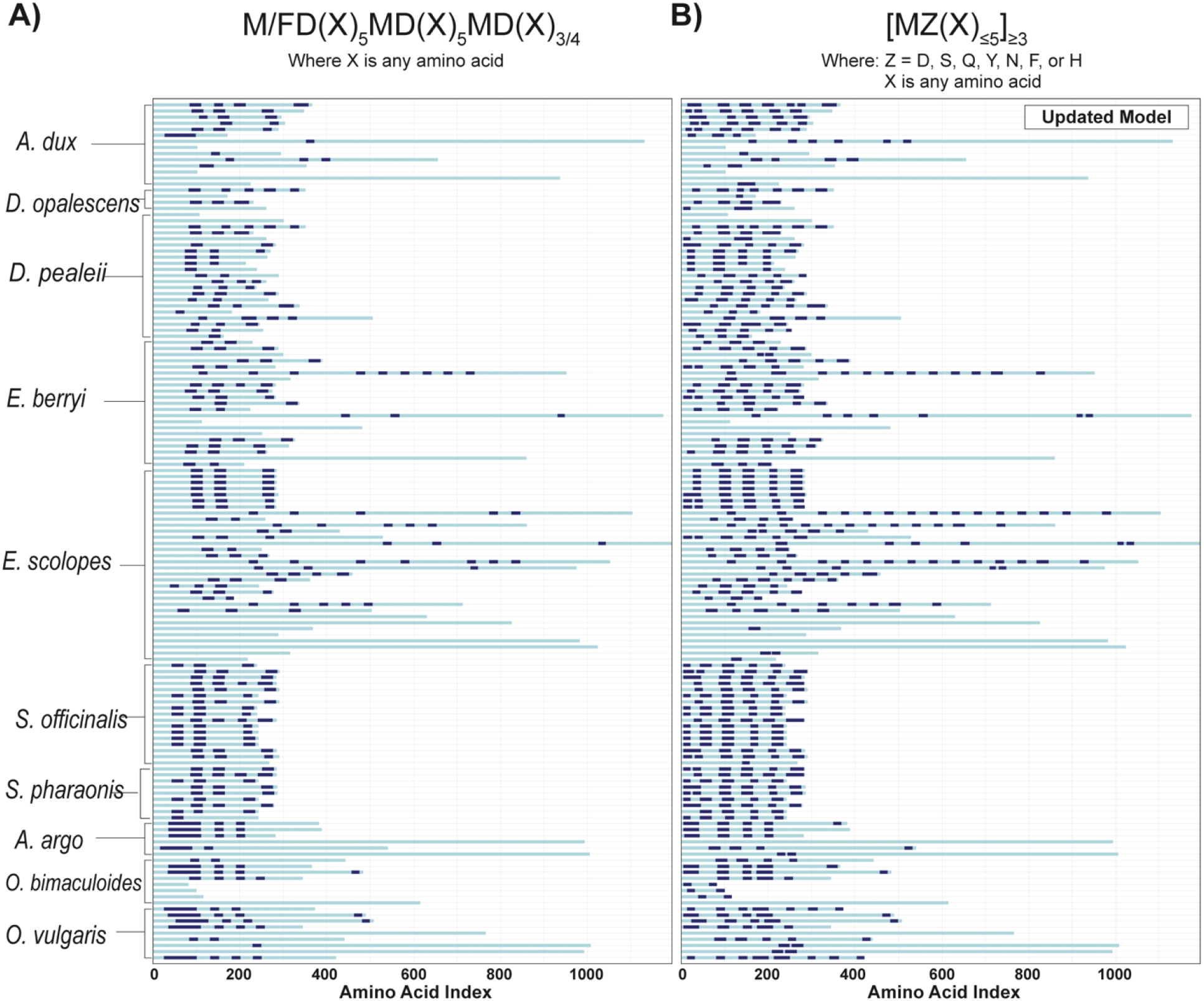
Domain-linker architecture of reflectin sequences under different domain definitions. A) Domains identified using the original definition M/FD(X)_5_MD(X)_5_MD(X)_3/4_. B) Domains identified using our simplified definition with relaxed Z-parameter constraints. Dark blue indicates domains, whereas light blue indicates linkers.

Based on these criteria, we simplified the reflectin domain as [MZ(X)_<5_]_>3_. This relaxed definition enabled us to identify 560 repeating sequences, or domains, from the 141 cephalopod reflectin sequences. Of these 560 identified domains, 53.6% contained three methionine repeats, whereas the remaining 46.4% contained four or more methionine repeats (**Figure 2B, Figure S1**). When comparing the distribution of methionine repeats across reflectin domain by species, cuttlefish including *S. officinalis* and *S. pharaonis*, exhibited a higher abundance of four-repeat domains compared to three-repeat domains. This sequence-level analysis captured the expansion of identified reflectin domains with increased domain coverage from 19.3% to 31.8% without compromising specificity, as demonstrated by negative control analyses showing no increase in false domain identification (**Figure S2)**. This improvement indicates that the original constraints likely excluded functionally relevant domains in other species.

Expanding on these observations, we created a matrix of sequence alignments using a Biopython toolset.^53^ Using the PairwiseAligner method with the ‘BLASTp’ scoring function,^54^ each domain in the reflectin sequence was aligned with every other domain across the dataset. Then, for each sequence, every residue was scored based on whether its reflectin domain parity agreed with that of the aligned residue on each other sequence. Parity agreements between residues were assigned a score of 1.0, alignment gaps were assigned a score of 0.0, and parity disagreements were assigned a score of -1.0, with the total sequence-sequence relationship represented by the average of all point scores for the target sequence (hereafter referred to as the Reflectin Domain Alignment Score or RDAS). The RDAS for the total dataset was then visualized as a heatmap (**Figure 3**), revealing trends in reflectin domain placement and conservation patterns both within and between species to be visually observed. Comparisons across the RDAS patterns using the original parameters (**Figure 3A)** versus our relaxed Z-position tolerance parameters (**Figure 3B**) revealed that the original parameters produced compressed similarity distributions with predominantly moderate scores. In contrast, high-similarity domain alignment pairs emerged more clearly using the expanded Z-position tolerances, producing more interpretable species-level clustering. Given this visual comparison, 3 notable features emerge from our relaxed definition. The first is that octopus species appear to have the lowest domain alignment, suggesting more variability in these proteins. The second is that cuttlefish species appear to have the highest domain alignment not only within itself but across all species. Third, the squid appear somewhere in the middle. Even though this analysis is surface level, it highlights focus areas to be investigated further in tying in sequence structures to functions. Furthermore, it would be interesting to explore where these reflectins are located within the animals. For instance, are sequences with higher average RDAS localized within a specific organ common to all animals? Likewise, could sequences with the lowest average RDAS indicate species-specific functions?

**Figure 3.**
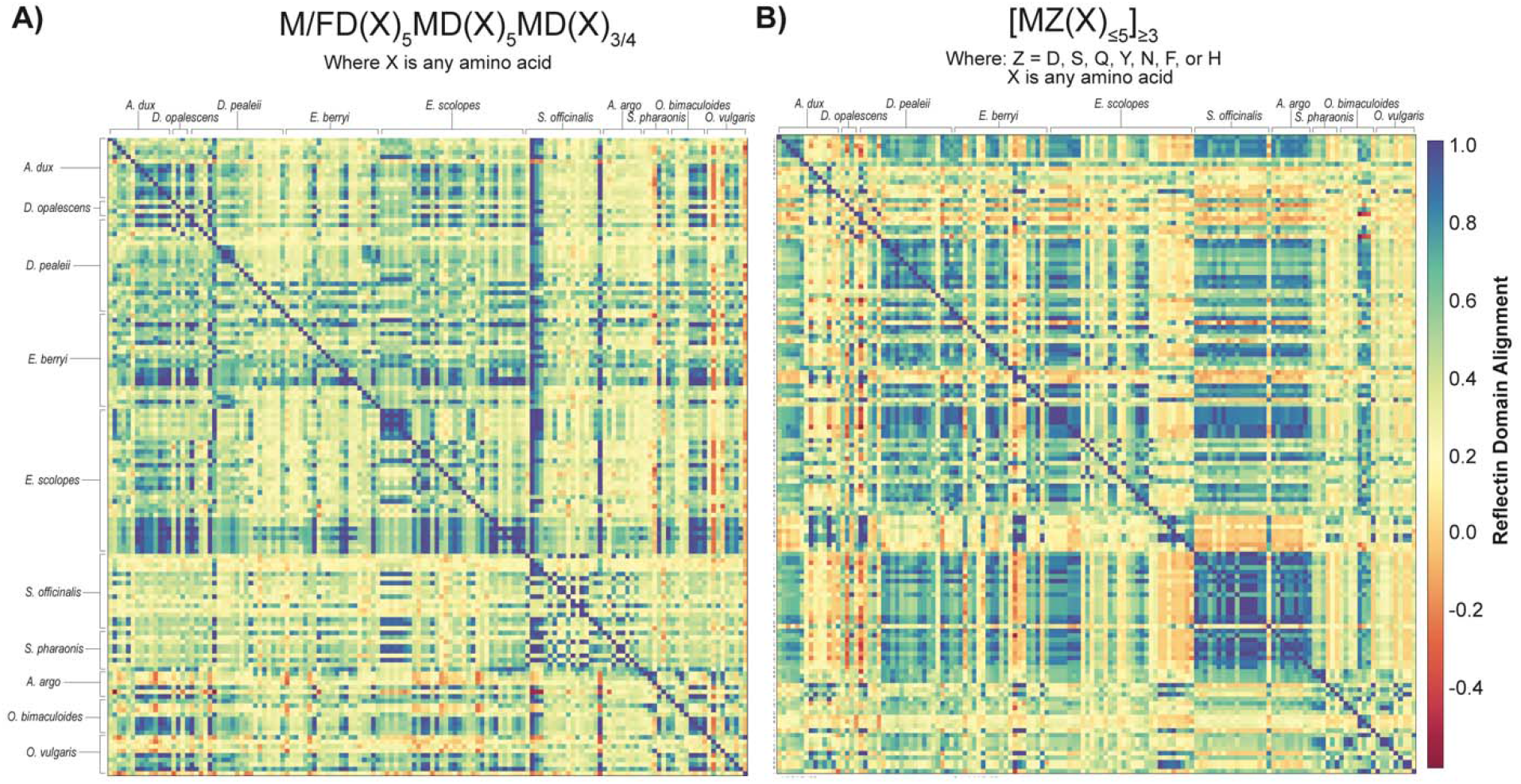
Heat map generation of reflectin domain alignment scores (RDAS) between and within cephalopod species. A) Alignment scores using the original reflectin defintion and B) alignment scores using our relaxed Z-parameter definition. The color gradient indicates reflectin alignment quality ranging from red (negative scores are dissimilar) to blue (positive scores are similar).

Given this dramatic increase in domain identification, rigorous validation of our expanded Z-position parameters was essential to ensure biological relevance and exclude the possibility of our result being caused by a computational artifact. To determine the functional consequences of reflectin domains, it is important to consider not only which amino acids are present but how they are distributed within conserved motifs. We therefore systematically analyzed the Z residue of the [MZ(X)_5_] repeating motif across all identified reflectin domains. From 2,000 Z-position residues across 141 reflectin sequences, we determined the frequency distribution of amino acids occupying this critical position (**Figure S3**). To establish principled boundaries for Z-position tolerance, we measured the total possibility space by counting all post-methionine amino acids in the complete reflectin dataset (**Figure S3**). Based on this density distribution, we constructed 20 progressive models, each incrementally expanding the set of valid Z residues. For each model, domain coverage (C) was measured as a percentage of total sequence length for both reflectin sequence and negative controls (**Figures S4A-B)**. Specificity was quantified using the metric ΔC_reflectin_/ΔC_negative_, where values greater than 1.0 indicate preferential domain identification in reflectin sequences over negative controls (**Figure S4C)**. Residues that increased reflectin domain identification without sacrificing specificity (ΔC_reflectin_/ΔC_negative_ > 100) while being sufficiently represented in Z positions (>1 occurrence per reflectin sequence on average) were selected for the expanded model. Based on these stringent criteria, an expanded model which accommodates this new, larger set of reflectin sequences was found to be one in which the Z-position residue could be: aspartic acid, serine, glutamine, tyrosine, asparagine, phenylalanine, or histidine. This differs from the original reflectin definition, where Z was restricted to aspartic acid.

This expanded Z-position amino acid set provides functional diversity at a structurally critical location. Positioned immediately after the methionine, but before the X spacer the Z-position may act as a ‘molecular control switch’ that regulates how tightly or loosely reflectin domains can bind together. The inclusion of charged (aspartic acid), polar (serine, glutamine, tyrosine, and asparagine), and aromatic (phenylalanine, tyrosine, and histidine) residues enable multiple interaction modes such as electrostatic networks, hydrogen bonding, and π-π stacking that could fine-tune both assembly stability and optical properties. This positional and chemical diversity may explain why extended domains with varied Z-residues are particularly abundant in species like *Doryteuthis*, where rapid, precise control over reflectin assembly is essential for signaling.

It is important to note that every domain is positioned adjacent to at least one cationic linker;^52^ however, little has been done beyond the original characterization to understand the functional implications of these linkers. To investigate this, we examined amino acid composition patterns across all linkers (i.e., any sequence in the protein that does not fit our new reflectin domain definition). We observed that linker regions showed distinct compositional preference for tyrosine (12.0%), asparagine (8.7%), arginine (8.5%), glycine (7.9%), and proline (7.3%) (**Figure S5)**. We suspect the elevated tyrosine content could facilitate interdomain aromatic interactions which are critical for higher-order assembly, while proline enrichment could provide structural kinks due to its backbone rigidity. The hydrogen bonding capacity of asparagine could aid in modulating domain spacing, and the continued presence of arginine could enable charge-based interactions between domains and linkers that coordinate assembly responses.

Beyond overall linker composition, we investigated whether specific positional constraints existed within linker regions relative to adjacent domains. To test this, sequence logos were constructed to examine trends in linker identity using a set of continuous strings comprising all amino acids outside of reflectin domains. Linker sequences between domains were extracted, excluding terminal regions, trimmed to 10 positions, and submitted to the Seq2LogoServer for logo generation using a Kullback-Leibler logotype (**Figure 4**).^55^ This analysis revealed positional specificity within the critical near-domain region: at position 2 from the domain boundary, asparagine (56.0%) and aspartic acid (21.1%) dominate, providing amine functionality and negative charge; position 3 shows overwhelming proline preference (87.5%); and position 4 is enriched for aromatic residues phenylalanine (50.9%) and tyrosine (21.0%). This precise but limited positional organization suggests that only the immediate domain-adjacent region positions (2-4) are functionally constrained, while the remainder of linker regions remain compositionally flexible. Beyond position 4, the lack of positional constraints may allow for species-specific sequence variation while maintaining the core functional architecture necessary for coordinated domain assembly for structural iridescence.

**Figure 4.**
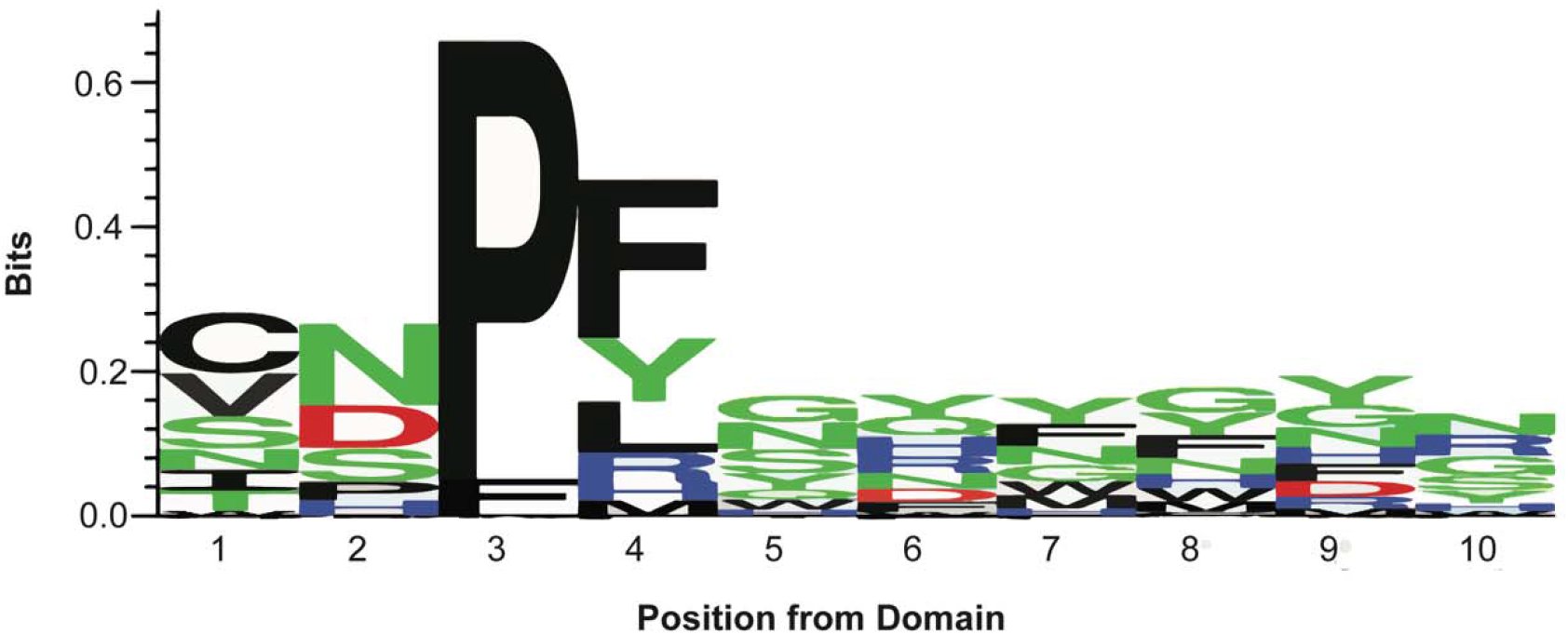
Sequence logo showing amino acid conservation in positions following a domain under the updated domain definition. Residue height (in bits) represents relative enrichment. Position indicates distance from the domain boundary.

## CONCLUSIONS

Our analysis introduces a redefined framework that relaxes the reflectin domain definition to include extended repeat sequences. This expansion was essential to capture the full diversity of newly available reflectin sequences across multiple cephalopods. While our computational analysis reveals obvious regions of strong similarity across sequences and species, stronger connections to function still need to be established through experimental validation. Key experiments could include point mutations in recombinant expression or in situ hybridization studies evaluating the location and sequestration of the sequences during development.

The domain architecture is the signature feature that differentiates reflectins from all other self-assembling proteins. The exceptionally high methionine content (22.4%) provides a hydrophobic anchor,^56^ while the abundance of charged residues creates extensive electrostatic networks that enable rapid, reversible assembly changes during optical switching.^57-64^ Interestingly, the two identified species that possess stimuli-responsive iridophores, *D. opalescens* and *D. pealeii*, are the only species in our dataset to possess reflectins with up to seven or eight methionine repeats (**Figure S1**). Yet, our analysis with RDAS reveals previously unidentified sequence similarities across other species, which may point to other adaptive reflectins. These sequence level comparisons could also inform the rational design of other proteins, beyond reflectins, that could be directly harnessed when developing next generation photonic materials. Furthermore, as additional genomic information becomes available, we may find that these trends change. Redefining criteria constituting what makes a reflectin domain or motif – and by extension what defines a reflectin protein – is important to adapt as more information becomes available. For now, our new framework for identifying and classifying these sequences reveals connections between domain and linker architectures that should be considered when interrogating the full diversity of cephalopod-based or inspired photonic systems.

## ACKNOWLEDGMENTS

This work was supported by the Air Force Office of Scientific Research (FA9550-22-1-0467) and the Marine Biological Laboratory Early Career Fellows gift from Susan and David Hibbitt.

## DATA AVAILABILITY

The data that support this work are available within the article and its Supporting Information. The automated assignment of reflectin domains and its visualization were created using an in-house script available at https://github.com/LeilaDeravi/Reflectin1.git.

## METHODS

### Mining uncharacterized reflectin sequences

The *Doryteuthis pealeii* Reflectin-A1 protein sequence was used as bait to search the proteomes of *E. berryi, E. scolopes, D. pealeii, Octopus bimaculoides, Architeuthis dux, Argonauta argo, and Octopus vulgaris* using BLAST NCBI. Additional reflectin sequences from previous studies were also added.

### Automated assignment of reflectin domains

The full sequence set was imported, and each sequence was analyzed iteratively for the presence of reflectin domains by selecting a test sequence and using the methionine initiation pattern as a critical marker. After identifying all methionine residues in the test sequence, the Z position residue of each methionine was evaluated for validity as a domain pattern initiator. This filtered subset of pattern initiators was then sorted based on the sequential distance to the next pattern initiator (a proxy for the number of X residues in the pattern) according to a set of heuristics:

1. If the number of X residues was at least one, but no more than five, the pattern had a ‘Valid’ length.
2. If the number of X residues was zero (i.e., there were two initiator patterns in direct succession: MZMZ), a ‘grace frame’ of two residues was added to the initiator pattern for the purpose of length determination. If this did not yield a ‘Valid’ length, the pattern had a ‘Short’ length and was discarded.
3. If the number of X residues was greater than five, the pattern had a ‘Long’ length and indicated a break in a reflectin domain, either as the trailing end of a complete reflectin domain or as an invalidating abridgement.

The sequence of patterns was then evaluated for the presence of continuous chains of ‘Valid’ length patterns (which must, by definition, end in a single pattern of ‘Long’ length). A chain of at least three patterns was considered to constitute a reflectin domain. These data were then aggregated and applied at the sequence level, where each residue is labeled as being either ‘In’ or ‘Out’ of a reflectin domain. This process was performed for each sequence in the set. The assignment method is modular, accepts any number of sequences, and was used both for the reflectin database and our negative control set.

### Construction of a negative control data set

A negative control sequence set was constructed for the purpose of comparison to that of the known reflectins. All protein sequences labeled under the class Cephalopoda in the UniProt database were accessed.^65^ To reduce processing strain and data management concerns, 5,000 sequences were selected at random to make up the negative control sequence set. The protein sequences contained within this set were expressed in similar organisms as those in the reflectin set but were not expected to display reflectin activity.

