## Supplemental Information for "Cephalopod Genome Expansion Drives Broader Reflectin Domain Boundaries"

**Table S1.** Known and uncharacterized reflectin protein sequences retrieved from Uniprot and the Marine Biological Laboratory. Database source, accession numbers and species of origin are provided for each sequence.

|  | **Database Source:** | **Entry**  **name:** | **Species:** | **Available Protein Sequence:** |
| --- | --- | --- | --- | --- |
| 1 | “Reflectin-like protein A1” from UniProt | A0A088MEP7 | *Doryteuthis opalescens* | MNRYLNRQRLYNMYRNKYRGVMEPMSRMTMDFQGRYMDSQGRMVDPRYYDHYGRMHDYDRYYGRSMFNQGHSMDSQRYGGWMDNPERYMDMSGYQMDMQGRWMDAQGRYNNPFSQMWHSRQGHYPGYMSHHSMYGRNMHYPYHSHSASRHFDSPERWMDMSGYQMDMQGRWMDNYGRYVNPFHHHMYGRNMFYPYGSHCNNRHMEHPERYMDMSGYQMDMQGRWMDTHGRHCNPLGQMWHNRHGYYPGHPHGRNMFQPERWMDMSSYQMDMQGRWMDNYGRYVNPFSHNYGRHMNYPGGHYNYHHGRYMNHPERQMDMSGYQMDMHGRWMDNQGRYIDNFDRNYYDYHMY |
| 2 | “Reflectin-like protein C1” from Uniprot | A0A088MRL7 | *Doryteuthis opalescens* | MNKHSSSHGMHGENYSRTGARGLHRGMEHESKSMYGKGRERSTDHGDMESSRHGMPGGMNPGMYGGMPSGMPGGMYPGMYGGMPGFGVPQMMQCPPDMPRRYIDRHDRSMDMPYGRYMDMPQGRYMSSQDRLMHMMHNRHLYGRMMDQGRMGEPMEGNMENRGRNMENYE |
| 3 | “Reflectin-like protein A2” from Uniprot | A0A088MI16 | *Doryteuthis opalescens* | MNRYMMRHRPMYSNMYRTGRKYRGVMEPMSRMTMDFQGRYMDSQGRMVDPRYYEYGRCHDYDRYNGRSMFNNGPYMDGQRYGGWMDFPERYMDMSGYQMDMHGRWMDSQGRYCNPMGHSWSNRQGYYPGSNYGRNMFNPERYMDMSGYQMDMQGRWMDMGGRHVNPFSHSMYGRNMFNPSYFSNRHMDNPERYMDMSGYQMDMQGRWMDTQGRYMDPSMSNMYDNYNYWY |
| 4 | “Reflectin-like protein B1” from Uniprot | A0A088MEC8 | *Doryteuthis opalescens* | MSSFMDPMHYDGMGMSHSKSGDFSHNCMRSFHKSQRDGMRRDIMGKSSKNRRFGNLMEPMSRMTMDFHGRLIDSQGRIVDPGHYFAMDDHYMENDRFLYPHDMLRNRHGMYGFMQGDYGNNMHRGMFADGMYRDMHHSGMNPSSYMHGGSMQNRPMMYMQGRYLDDSYFMNYHDPPVIVHSHYNDQEGRHQGMYDRHSDSYGSHRRHGDSHSMPRRPSESHSPQRRPSEGHIIQVRPEGGSSRKTSRAQLFPDDKLTDSA |
| 5 | “Reflectin-like protein A1” from Uniprot | D3UA43 | *Doryteuthis pealeii* | MNRYLNRQRLYNMYRNKYRGVMEPMSRMTMDFQGRYMDSQGRMVDPRYYDYYGRMHDHDRYYGRSMFNQGHSMDSQRYGGWMDNPERYMDMSGYQMDMQGRWMDAQGRFNNPFGQMWHGRQGHYPGYMSSHSMYGRNMYNPYHSHYASRHFDSPERWMDMSGYQMDMQGRWMDNYGRYVNPFNHHMYGRNMCYPYGNHYYNRHMEHPERYMDMSGYQMDMQGRWMDTHGRHCNPFGQMWHNRHGYYPGHPHGRNMFQPERWMDMSGYQMDMQGRWMDNYGRYVNPFSHNYGRHMNYPGGHYNYHHGRYMNHPERHMDMSSYQMDMHGRWMDNQGRYIDNFDRNYYDYHMY |
| 6 | “Reflectin-like protein A2” from Uniprot | D3UA44 (also known as Dopeav2126554m_Dpe41_22893158_22899332__110071_NA) | *Doryteuthis pealeii* | MNRYMMRHRPMYSNMYRTGRKYRGVMEPMSRMTMDFQGRYMDSQGRMVDPRYYDYGRCHDYDRYYGRSMFNYGPNMDGQRYGGWMDFPERYMDMSGYQMDMHGRWMDSQGRYCNPMGHSWSNRQGYYPGSNYGRNMFNPERYMDMSGYQMDMQGRWMDMGGRHVNPFSHSMYGRNMFNPSYFSNRHMDNPERYMDMSGYQMDMQGRWMDTQGRYMDPSWSNMYDNYNSWY |
| 7 | “Reflectin-like protein B1” from Uniprot | D3UA45 | *Doryteuthis pealeii* | MSSFMDPMHYDGMGMSHSKTGDFSHNCMRSFHKSQRDVMRRDIMGKSSKNRRFGNLMEPMSRMTMDFHGRLIDSQGRIVDPGHYFAMDDHYMENDRFLYPHDMLRNRHGMYGFMQGDYGNNMHRGMFADGMYRDMHHSGMNPSGYMHGGSMQNRPMMYMQGRYLDDSYFMNYHDPPVIVHSHYNDQEGRHHGMYDRHSDSYGSHRRHGDSHSMPRRPSESHSPQRRPSEGHIIQVRPEGGSSRKTSRAQLFPDDKLTDSA |
| 8 | Provided from MBL using Ref^1^ | Dopeav2089456m_Dpe26_91722086_91722938__110071_NA | *Doryteuthis pealeii* | MNRYMNRFRNWYGNNYRGRYRGMMEPMSRMTMDFQGRYMDSCGRMVDPRYYDYYGRWYDYDRYYGRSMFNYGWMMNGDRYNNYYRWMDFPERYMDMSCYQMDMYGRWMDMYGRQCNPFRQWWYYRHGYFPGYYYGCNMFYPERWIDMSNYYMDMQGRYMDRWGRYCNPFSHYYNYWNRYCNYPGYYNYYYMYYPERYFDMSNWQMDMQGRWMDMHGRYCNPYWYNWYGRHMYYPYQNYYWYGRWDYPWMDYSNWQMDMQGRWMDMQGRYMDFPYYYYYYNWY |
| 9 | Provided from MBL using Ref^1^ | Dopeav2015112m_Dpe04_109965726_109966538__110071_NA | *Doryteuthis pealeii* | MNRGSNRGMMEPMSRMTMDFQGRYMDSMGRMVDPRFNDYYGRWNDYDRYYGRSMFNYGWMMNGDRYNRNFRCMDFPERYMDMSGYQMDMCGRWMDSYGRQCNPFNQWSYNRHGYYPGYSYGRNMCYPERWMDMSNYCMDMQGRYMDRSGRHCNPFSQHMNYYGRYWNYPGYNNYYNRNMYYPERHFDMSNWQMDMQGRWMDRQGRYNNPYWCNWYGRNMYNPYQNNQWYGRYDYPGMDCSNWEMDRQGRGMDAQDHYMNSWMGDSCYNNW |
| 10 | Provided from MBL using Ref^1^ | Dopeav2015121m_Dpe04_110178292_110179080__110071_NA | *Doryteuthis pealeii* | MNRRRYRGVMEPMSRMSMDFQGRYMDSMGRMVDPRFNDYYGRWNDYDRYYGRSMFNYGWMMNGDRYNRNFRSMDFPERYMDMSGYQMDMCGRWMDSYGRQCNPFNQWSDNRDGYYPGYSYGRNTCYPERWMDMSNYCMDMQGRYMDRSGRHCNPFSQHMNYYGRYWNYPGYNNYYNRNMYYPERHFDMSNWQMDMQGRWMDRQGRYNNPYWCNWYGRNMYNPYQNNQSYGRWDHPGMDYSNWEGRGMDMQGDYMNSWVDDSC |
| 11 | Provided from MBL using Ref^1^ | Dopeav2015117m_Dpe04_110029247_110029885__110071_NA | *Doryteuthis pealeii* | MNRGSFRGMMEPMSRMSMDFQGRYMDSMGRMVDPRFYDYYGRWNDYDRYYGRSMFNYGWMMNGDRYNRNFRSMDFPERYMDMSGYQMDMCGRWMDSYGRQCNPFNQWSYSRHGYYPGYSYGRNMCYPERWMDMSNYCMDMQGRYMDRSGRHCNPFSQHMNYYGRYWNYPGYNNYYNRNMYYPERHFDMSNWQMDMQGRWMDRQGRYNNPTGVT |
| 12 | Provided from MBL using Ref^1^ | Dopeav2015124m_Dpe04_110194538_110195251__110071_NA | *Doryteuthis pealeii* | MNRGSYRGMMEPMSRMSMDFQGRYMDSMGRMVDPRFNDYYGRWNDYDRYYGRSMFNYGWMMNGDRYNRNFRCMDFPERYMDMSGYQMDMCGRWMDSYGRQCNPFNQWSYNRHGYYPGYSYGRNTCYPERWMDMSNYCMDMQGRYLDRSGRHCNPFSQHMNYYGRYWNYPGYNNYYNRNMYYPERHFDMSNWQMDMQGRWMDRQGRYNNPYWCNWYGRNMYNPYQNNQWYGRYDYPGMD |
| 13 | Provided from MBL using Ref^1^ | Dopeav2015120m_Dpe04_110134210_110135076__110071_NA | *Doryteuthis pealeii* | MNRCMNRYRPNMWGNRSNNNMWGNMNRRRYRGVMEPMSRMSMDFQGRYMDSMGRMVDPRFNDYYGRWNDYDRYYGRSMFNYGWMMNGDRYNRNFRCMDFPERYMDMSGYQMDMCGRWMDSYGRQCNPFNQWSYNRHGYYPGYSYGRNMCYPERWMDMSNYCMDMQGRYMDRSGRHCNPFSQHMNYYGRYWNYPGYNNYYNRNMYYPERHFDMSNWQMDMQGRWMDRQGRYNNPYWCNWYGRNMYNPYQNNQWYGRYDYPGMDCSNWEGRGMDMQGDYMNPWMDDSCYNN |
| 14 | Provided from MBL using Ref^1^ | Dopeav2015119m_Dpe04_110117727_110125075__110071_NA | *Doryteuthis pealeii* | MNDPCYKNMFYNMNNATTAVYEIIMNRCMNRYQPNNMWGNMSNNNMWDNNMWGNTFNNNNMSCNRGRYRGMMEPMSRMTMDFQGRYMDSMGRMVDPRFNDYYGRWNDYDRYYGRSMFNYDWMMNGDRYNRNFRSMDFPERYMDMSGYQMDMCGRWMDSYGRQCNPFNQWSYNRHGYYPGYSYGRNMCYPERWMDMSNYCMDMQGRYMDRSGRHCNPFSQYGRWDYPGMDGSNWDMDRQGRGMDMQDDYMNSWMGDSCYNN |
| 15 | Provided from MBL using Ref^1^ | Dopeav2126548m_Dpe41_22697540_22805831__110071_NA | *Doryteuthis pealeii* | MNRYMNRYRNMFNNNNMWGNMYRGRYRGMMEPMSRMTMDFQGRYMDSCGRMVDPRFYDYYGRWNDYDRYYGRSMFNYGWMMNGDRYNNYYRWMDFPERYMDMSGYQMDMYGRWMNPYGRQCNPFNQWSYNRHGYYPGYSYGRNMCYPERWMDMSNYSMDMQGRYMDRWGRQCNPFSQYMNWYGRYWNYPGYNNYYNRNMYYPERHFDMSNWQMDMQGRWMDMQGRYMDFPYYYYNWY |
| 16 | Provided from MBL using Ref^1^ | Dopeav2126408m_Dpe41_18680479_18729544__110071_NA | *Doryteuthis pealeii* | MNRYMNRYRPMFNNMWGNMYRGRYRGMMEPMSRMTMDFQGRYMDSMGRMVDPRYYDYYGRWNDYDRYYGRSMFNYGWMMNGDRYNNYYRWMDFPERYMDMSGYQMDMNGRWMDMYGRQCNPFNQWNYYRHGYYPGYSYGRNMFYPERWMDMSNYCMDMQGRYMDRWGRHCNPFSQYMNWYGRYWNYPGYNNYYYNRYMYYPERYFDMSNWQMDMQGRWMDMQGRYCNPYWYNWYGRHMYYPYQNYWYGRWDYPGMDCSNWQMDMQGRWMDMQGRYMDPCWWNDSYSYYY |
| 17 | Provided from MBL using Ref^1^ | Dopeav2126549m_Dpe41_22699478_22700272__110071_NA | *Doryteuthis pealeii* | MSNMWGNMFRGRYRGMMEPMSRMTMDFQGRYMDSCGRMVDPRFNDYYGRWNDYDRYYGRSMFNYGWMMNGDRYNNYYRWMDFPERYMDMSGYQMDMYGRWMNPYGRQCNPFNQWNYNRHGYYPGYSYGRNMCYPERWMDMSNYSMDMQGRYMDRWGRHCNPFSQHMNWYGRYWNYPGYNNYYNRNMYYPERHFDMSNWQMDMQGRWMDMQGRYNNPYWYNWYGRNMYYPYQNQWYGRWDYPGMDCGMDMQGGYMNSWMGDSCYNN |
| 18 | Provided from MBL using Ref^1^ | Dopeav2126550m_Dpe41_22719023_22724506__110071_NA | *Doryteuthis pealeii* | MNRCMNRYRPSNMWGNMSNNNMWGNMSNNNMWGNNMWGNNMWGNMSSNNMWGNMNRGRYRGMMEPMSRMTMDFQGRYMDSCGRMVDPRFNDYYGRWNDYDRYYGRSMFNYGWMMNGDRYNRNYRCMDFPERYMDMSGYQMDMCGRWMDPYGRQCNPFNQWSYNRHGYYPGYSYGRNMCYPERWMDMSNYSMDMQGRYMDRWGRHCNPFSQHMNWYGRYWNYPGYNNYYNRNMYYPERHFDMSNWQMDMQGRWMDMQGRHNNPYWYNWYGRNMYNPCQNNQWYGRGDYPGMDCSNWQMDMQGRGMDMQGRGMDMQGRGMDMQGGYMNSWMGDSCYNNW |
| 19 | Provided from MBL using Ref^1^ | Dopeav2126544m_Dpe41_22633913_22637615__110071_NA | *Doryteuthis pealeii* | MDMSCYQMDMYGRWMDNYGRHCNPFNQWSYNRHGYYPGYSYGRNMCYPERWMDMSNYSMDMQGRYMDRWGRHCNPFSQYMNWYGRYWNYPGYSNYYYNRYMYYPERYFDMSNWQMDMQGRWMDMQGRYNNPYWYNWYGRNMYYPYQNQWYGRWDYPGMDCGMDMQGGYMNSWMGDYCYNNW |
| 20 | Provided from MBL using Ref^1^ | Dopeav2126411m_Dpe41_18801087_18802604__110071_NA | *Doryteuthis pealeii* | MNRLHDRRRFMAFCFPRGDNYRGVMEPMPNMGMDFQGRYFDGQGRQMDPKMLDFYRRYHFGGQNSPCYHYGGHNYNWSPYYMGYMKGNWYPRYGRQMNFPEKFMDLSNYQMDMQGRWMDMQGQFANPFNSNQLKQQNCALVAPHMHLNNYMHLNNYMRYMLRPERFMEMPAYQMDMYNRYFLPSSNNPLALCARQNVNGVHPYMFNNWRCMFFPERHMDMSNYQMDMQGRWMDMQGRHNNPFYQMCSGRHGFYPGYGYGRNMFQPERWMDMSGYQMDQQGRWMDMCGRQVNPFGRNMFYPNRQMEYPERYMDMSGYQMDMQGRWMDGHGHNCNPLNQFGYNRQGFYPGYQQSRGMFNPERGMDMPGFQNDMQSRWMEMCGRHGRQMNHPAANYQYYARYMNYPERNMETPGYQGDLQGRWMDMCSRYMNQAYGYYGQRSAMGDDFGRHMGRPQQYLGMGNYMDMQGPHMRNTDYKNHHHNFYGCHQHPQMFNHHGYGNYAAKYDY |
| 21 | Provided from MBL using Ref^1^ | Dopeav2126555m_Dpe41_22909843_22910684__110071_NA | *Doryteuthis pealeii* | MNRYMNRFRNWYGNNYRGRYRGMMEPMSRMTMDFQGRYMDSCGRMVDPRYYDYYGRWNDYDRYYGRSMFNYGWMMNGDRYNNYYRWMDFPERYMDMSCYQMDMCGRWMNMYGRQCNPFNQWWYNRHGYFPGYYYGCNMYYPERWMDMSNYYMDMQGRYMDRYFDMSNWQMDMHGRWMDMHGRYNNPYWYNWYGRHMYYPYQNYYWYGRWDYPGMDYSNWQMDMQGRWMDMQGRYMDFPYYYYNWY |
| 22 | Provided from MBL using Ref^1^ | Dopeav2126542m_Dpe41_22614764_22615522__110071_NA | *Doryteuthis pealeii* | MWGNMYRGRYRGMMEPMSRMTMDFQGRYMDSCGRMVDPRFYDYYGRWNDYDRYYGRSMFNYGWMMNGDRYNNYYRWMDYPERYMDMSCYQMDMYGRWMDNYGRHCNPFNQWSYNRHGYYPGYSYGRNMCYPERWMDMSNYSMDMQGRYMDRWGRHCNPFSQYMNWYGRYWNYPGYNNYYYNRNMYYPERYFDMSNWQMDMQGRWMDMQGRYNNPYWYNWYGRNMYYPYQNQWYGRWDYPGMDCGMDMQGGYMN |
| 23 | Provided from MBL using Ref^1^ | Dopeav2126545m_Dpe41_22657072_22683550__110071_NA | *Doryteuthis pealeii* | MFDMNWYGRYWNYPGYSNYYYNRYMYYPERYFDMSNWQMDMQGRWMDMQGRYNNPYWYNFPSLRHTHFICYDEATDWDFPERYMDMSGYQMDMYGRWMNMYGRQCNPFNQWNYNRHGYYPGYSYGRNMFYPERWMDMSNYCMDMQGRYMDRWGRHCNPSPST |
| 24 | Provided from MBL using Ref^1^ | >Dopeav2126409m_Dpe41_18775466_18783571__110071_NA | *Doryteutheis pealeii* | MRHFNNPRPFMGLMEPMSRMTMDFEGRFVDSRGRLVDPLHYRPRQYRAWYDMLSNRHQDQSDVYRFESNSYGSQLDDCDRFWPLDDYESRLELYTGMNLDFHDNFEC |
| 25 | Provided from MBL using Ref^1^ | Dopeav2126420m_Dpe41_19019522_19050447__110071_NA | *Doryteutheis pealeii* | MSRQVKTSQGNTTGPKNTAPQSSQSIPHTSQSATKSTHSMPQSAQSMPHGSQSLPRGPPSMPRGPQSMHDAYSMPYGGYDAYSMPPHGTYPMGYGAPPPMSHGGFPMSHDPQSMPRGMYPMYGGGGPVSSSMSQEENKNRMKTFSKSQRDTLSREGVKSGSSHQRGSVPESMNRMTMDYQGRFIDSRGRVVDYSHNMDSSDGLHDRRYMFARDMPREEQVMYDYRMPEYMYGNMMENPDMMMDMEMQGRYMPFYDQYADDDYNDGILDYMDGYNSDFPGMHFPESRTMDSAGRYGDSFER |
| 26 | “Reflectin 2b” from Uniprot | Q6WDN4 | *Euprymna scolopes* | MNRYMNRFRNFYGNMYRGRYRGMMEPMSRMTMDFQGRYMDSQGRMVDPRFYDYYGRYNDYDRYYGRSMFNYGWMMDGDRYNRCNRWMDYPERYMDMSGYQMDMYGRWMDMQGRHCNPYSQWMMYNYNRHGYYPNYSYGRHMFYPERWMDMSNYSMDMYGRYMDRWGRYCNPFYQFYNHWNRYGNYPGYYNYYYMYYPERYFGMSNWQMDMQGRWMDMQGRYCSPYWYNWYGRHMYYPYQNYYWYGRYDYPGMDYSNWQMDMQGRWMDMQGRYMDYPYNYYNWNH |
| 27 | “Reflectin 2c”from Uniprot | Q6WDN3 | *Euprymna scolopes* | MNRYMNRFRNFYGNMYRGRYRGMMEPMSRMTMDFQGRYMDSQGRMVDPRYYDYYGRYNDYDRYYGRSMFNYGWMMDGDRYNRYNRWMDFPERYMDMSGYQMDMYGRWMDMQGRHCNPYSQWMMYNYNRHGYYPNYSYGRHMFYPERWMDMSNYSMDMYGRYMDRWGRYCNPFYQFYNHWNRYGNYPGYYNYYYMYYPERYFDMSNWQMDMQGRWMDMQGRYCSPYWYNWYGRQMYYPYQNYYWYGRYDYPGMDYSNYQMDMQGRYMDMQGRYMDYPYNYYNWNH |
| 28 | “Reflectin 2d” from Uniprot | Q6WDN2 | *Euprymna scolopes* | MNRYMNRFRNFYGNMYRGRYRGMMEPMSRMTMDFQGRYMDSQGRMVDPRYYDYYGRFNDYDRYYGRSMFNYGWMMDGDRYNRYNRWMDFPERYMDMSGYQMDMYGRWMDMQGRHCNPYSQWMMYNYNRHGYYPNYSYGRHMFYPERWMDMSNYSMDMYGRYMDRWGRYCNPFYQFYNHWNRYGNYPGYYSYYYMYYPERYFDMSNWQMDMQGRWMDMQGRYCSPYWYNWYGRHMYYPYQNYYWYGRYDYPGMDYSNWQMDMQGRWMDMQGRYMDYPYNYYNWNH |
| 29 | “Reflectin 2a” from Uniprot | Q6WDN5 | *Euprymna scolopes* | MNRYMTRFRNFYGNMYRGRYRGMMEPMSRMTMDFQGRYMDSQGRMVDPRYYDYYGRYNDYDRYYGRSMFNYGWMMDGDRYNRYNRWMDFPERYMDMSGYQMDMYGRWMDMQGRHCNPYSQWMMYNYNRHGYYPNYSYGRHMFYPERWMDMSNYSMDMYGRYMDRWGRYCNPFYQFYNHWNRYGNYPGYYNYYYMYYPERYFDMSNWQMDMQGRWMDMQGRYCSPYWYNWYGRHMYYPYQNYYWYGRYDYPGMDYSNWQMDMQGRWMDMQGRYMDYPYNYYNWY |
| 30 | “Reflectin 3a” from Uniprot | Q6WDN6 | *Euprymna scolopes* | MNRYMNRFRNFYGNMCRNRNRGMMEPMSRMTMDFQGRYMDSQGRMVDPRYYDYYGRYNDYDRYYGRSMFNYGWMMDGDRYNRYNRWMDYPERYMDMSGYQMDMYGRWMDMQGRHCNPYSQWMMYNYNRHGYYPNYSYGRHMFYPERWMDMSNYSMDMYGRYMDRWGRYCNPFYHYYNHWNRSGNNPGYYSYYYMYYPERYFDMSNWQMDMQGRWMDMQGRYCSPYWYNWYGRQMYYPYQNYYWYGRWDYPGMDYSNWQMDMQGRWMDMQGRYMDPWWMNDSYYNNYYN |
| 31 | “Reflectin 1b” from Uniprot | Q6WDN7 | *Euprymna scolopes* | MNRFMNKYRPMFNNMYSNMYRGRNRGMMEPMSRMTMDFQGRYMDSQGRMVDPRYYDYYGRFNDYDRYYGRSMFNYGWMMDGDRYNRYNRWMDYPERYMDMSGYQMDMSGRWMDMQGRHCNPYSQWGYNYNRHGYYPNYSYGRHMFYPERWMDMSGYQMDMQGRYMDRWGRYCNPFSQYMNYYGRYWNYPGYNSYYNSRNMFYPERYFDMSNWQMDMQGRWMDNQGRYCSPYWNNWYGRQMYYPYQNNYFYGRYDYPGMDYSNYQMDMQGRYMDQYGMNDYCY |
| 32 | “Reflectin 1a” from Uniprot | Q6WDN8 | *Euprymna scolopes* | MNRFMNRYRPMFNNMYSNMYRGRYRGMMEPMSRMTMDFQGRYMDSQGRMVDPRYYDYYGRFNDYDRYYGRSMFNYGWMMDGDRYNRYNRWMDYPERYMDMSGYQMDMSGRWMDMQGRHCNPYSQWMMYNYNRHGYYPNYSYGRHMFYPERWMDMSNYSMDMYGRYMDRWGRYCNPFSQYMNYYGRYWNYPGYNNYYYSRNMYYPERYFDMSNWQMDMQGRWMDNQGRYCSPYWNNWYGRHMYYPYQNNYFYGRYDYPGMDYSNYQMDMQGRYMDQYGMNDYYY |
| 33 | Provided from MBL using Ref^2^ | Esc_cluster_11038p | *Euprymna scolopes* | MNQIPDRKRFMPHQFYKSEKYPGELEPITMMTMGFQGRYLDSQGRMHDPRVYESYRRYHIGGSNNLNSDPSCSQSYTWWPYPGDYTKPDWYPRPRRQMVFPEKFMDLSSYQMDMKGRWMDMQGRYTNPFNSKNRTRRNHFPLFSPQMYSNNYGNDMSHPEGNKDTPGYPMDSQDHWMYIQGRQTKPSSYNTVGFLGKQYPYKYPPYMHDTWRFISWPERYMDMTGYQMDMRGRWMDTQGRHCNPFNQCGYNKQGSYLGDPYNRNIVYSDKLMDRSNDRMDRQESSDGRYGRRVNPLSRHSCRVCMDMNNPYPGKMDMSNYQMDMQGRWMDTKGRYTNPFPSSGYNKQGPFHGFQHKHNMPYPEKLIDMSNYQMDMQGRWMDTQGRYTNPFSVCGYNRQVPFSGFPYNRNMHYPGKMIDMSNYQMDMQGRWMDTQGRYTNPFSFSSFTRQGPIPGFPYNRNMPYPEKMMDMSNYQMDMQGRWMDTQGRYANPFSLSSFTRQGLFPVFPYNRNMPYPEKMIDMSNYQMDMQGRWMDTQGRYTNPFSFGSFTRQGPIPGFPYNRNMPYPEKMIDMSNYQMDMQGRWMDTQGRYTNPFSFGSFTRQIPFPGFAYNRNMPYPEKMIDMSNYQMDMQGRWMDTQGRYTNPFSFGSFTRQGPIPGFPYNRNMPYPEKMIDMSNYQMDMQGRWMNTQGRYTNPFNFSSFTRQGPFPGFPYNRNMPYPEKMIDMSNYQMDMQGRWMDTQGRYTNPFSFSGYNKQELFPGFSYNRNIAYPENMMDMSNYQMDMDGHWMDTQGRYTNPFGFSSYKRQWPFSGLPYSCNMPYPENMMDMSNYQMDMQGRWMDMYGRYTNPFSFSGFNRRRPFHSFPYNRNMSYPENMIDMSNYQMDMQGRWMDTQGRYRNPFNQYSNYNRYGYFPEYSFDRNTFPMRFNNQIEAPGRFSQPYDHYLSTKDIVYPFYNYRYSHYIYYPDRYMDMSGNHVDMQEPSMDTESRFYPGYQQGRSMNPLNRLLDSHDYNPQTDMEGRWMDYYGEHTHPGFGDQERRTLGDYFRSLPGTYHGDTYRYLGMRNMYDRDQFHYQYGYYQQPPVINSHNDGYYDNNMMGYFYDN |
| 34 | Provided from MBL using Ref^2^ | Esc_cluster_10790p | *Euprymna scolopes* | SCTRTYLKLRQRYRSCPVAVKKHKNKQKKTATTKVAIMNRYMMKHRPMYNQMCRTGRRYRGVMEPMSRMTMDFQGRYMDSQGRMVDPRYYDFSGSSDRYSGRSMFNYGSYMDGGQRYGGFMDNPERYMDMSNYQMDMHGRWMDTQGRYNSPFSYYGYNRHGNYPSYYSYNRSMCNPERMMDMSNYQMDMQGRWMDNYGRHVNPFSHFMYGRNMHYPNFNYYSGRYMDYSDMSNPQMDMQGRYMDSSMSNMYDNYNNYY |
| 35 | Provided from MBL using Ref^2^ | Esc_cluster_11991p | *Euprymna scolopes* | MNQIPDRKRFMPHQFYKSEKYPGELEPITMMTMGFQGRYLDSQGRMHDPRVYESYRRYHIGGSNNLNSDPSCSQSYTWWPYPGDYTKPDWYPRPRRQMVFPEKFMDLSSYQMDMKGRWMDMQGRYTNPFNSKNRTRRNHFPLFSPQMYSNNYNKQGPFHGFQHKHNMPYPEKLIDMSNYQMDMQGRWMDTQGRYTNPFSVCGYNRQVPFSGFPYNRNMHYPGKMIDMSNYQMDMQGRWMDTQGRYTNPFSFSSFTRQGPIPGFPYNRNMPYPEKMMDMSNYQMDMQGRWMDTQGRYTNPFSVCGYNRQVPFSGFPYNRNMHYPGKMIDMSNYQMDMQGRWMDTQGRYTNPFSFSSFTRQGPIPGFPYNRNMPYPEKMMDMSNYQMDMQGRWMDTQGRYANPFSLSSFTRQGLFPVFPYNRNMPYPEKMIDMSNYQMDMQGRWMDTQGRYTNPFSFGSFTRQGPIPGFPYNRNMPYPEKMIDMSNYQMDMQGRWMDTQGRYTNPFSFSGYNKQELFPGFSYNRNIAYPENMMDMSNYQMDMDGHWMDTQGRYTNPFGFSSYRRQWPFSGLSYSCNMPYPENMMDMSNYQMDMQGRWMDMYGRYTNPFSFSGFNRRRPFHSFPYNRNMSYPENMMDMSNYQMDMQGRWMDTQGRYRNPFNQYSNYNRYGYFPEYSFDRNTFPMRFNNQIEVPGRFSQPYDHYLSTKDIVYPFYNYRYSHYIYYPDRYMDMSGNHVDMQEPSMDTESRFYPGYQQGRSMNPLNRLLDSHDYNPQTDMEGRWMDYYGEHTHPGFGDQERRTFGDYFRSLPGTYHGDTYRYLGMRNMYDRDQFHYQYGYYQQPPVINSHNDGYYDNNMMGYFYDN |
| 36 | Provided from MBL using Ref^2^ | Esc_cluster_13716p | *Euprymna scolopes* | MTAVGVIWIVTVSVVVVTPLVVRVAVASLHIHPASVHVHLVARHVHVPLGEVHPSVVPVVPITVHHPSVVKHGSSVVTVVVVESSVVVVVPGVDHSSLGVHVSSLEVHGHAGHRFHHTSVSTTVHVAVHVVEHGSVSVHVTIHGDLARMNRYMNRYRPMFNNMYGNMYRGRYRGMMEPMSRMTMDFQGRYMDSQGRMVDPRYYDYYGRFNDYDRYYGRSMFNYGWMMDGDRYNRYNRWMDFPERYMDMSGYQMDMYGRWMDMQGRYCNPYNQWGYNYNRHGYYPNYSYGRHMFYPERWMDMSGYQMDMQGRYMDRSGRYCNPFSQYMNYYGRFWNYPGYNSYYNRNMYYPERHFDMSNWQMDMQGRWMDNQGRYSSPYWNNFYGRQMYYPYQNYYSYGRYDYPGMDYSYSQMDGRFNDSWMGDSYYNNW |
| 37 | Provided from MBL using Ref^2^ | Esc_cluster_9945p | *Euprymna scolopes* | MNRFMNRYRPMFNNMYSNMYRGRYRGMMEPMSRMTMDLQGRYMDSQGRMVDPRYYDYYGRFNDYDRYYGRSMFNYGWMMDGDRYNRYNRWMDYPERYMDMSGYQMDMSGRWMDMQGRHCNPYSQWMMYNYNRHGYYPNYSYGRHMFYPERWMDMSNYSMDMYGRYMDRWGRYCNPFSQYMNYYGRYWNYPGYNNYYYSRNMYYPERYFDMSNWQMDMQGRWMDNQGRYCSPYWNNWYGRHMYYPYQNNYFYGRYDYPGMDYSNYQMDMQGRYMDQYGMNDYYYMSAIPVVPIGAAVASLVIHPASLHIHLPVRHIEVSFWVVHVSAVIIVVVPGVVPVSAIVIHVLGERVAVTSPPVHVTSVHVHGVVRHVHPSLRVEHMAAVGVVGIVTVSVVVVHHPLAIRVAVASLHVHPASRHVHLVARHVHVPLGVVHPSVVSVVPITVHHPAVVEHGSSVVTVIVVESSIVVVVSGVDHSSLGVHVSSLEVHGHTGHGFHHTSVSTAVHVAIHVVEHGSVSVHKTVHDDFLR |
| 38 | Provided from MBL using Ref^2^ | Esc_cluster_24511p | *Euprymna scolopes* | MRNQTKSGGFNKTSTLDTNTPGTSASSQPSLDHSDLKGKLPSLQHIDPDYRFRSNMIPKRHPMFPMNGPHRMMPPNARHANRMHSPNVMFRGRAPNMMPGRSMFQGPGAQMGMAPPGGFMAPMQHRPKESENIQPQNEGQGRMSGFTQRQEEMPKDSRGFPGKMMDFHTIPGQPKHGGPMRMHTGPMGMQSGPMGMRSGPMGMHSGPMGKQSGPMKMHSGPIGMHSGPMGMQSGPMGMQGGPMGKQSGPMGMHSGPIGVQSNPMGKQSGPMGMHGGPMRMQSGLMGMRGGPMGKQSGPMGMQSGPMGIHGPMGMQGGPMGILGSPMGKQSSPMRMHSGTMRMQSGPMRMRSGPMGMKGGPMGMQSGLMRMRSAPMGMHAFGMNKTSGQQGKPMGRFPRGLRRHMMPMPLSYKRMIRFPERYMDLSGYTMDFKGRFVNQQGKHVNPLHSARMHQGRAMMPAPLPIKRRFMQKPERYMDLSRYTMDFEGRYRDRYGRQIDPMEQFNANIGRYVRNLPPVHRRMVRFPERYMDLSRYTMDFQGRFMDSHGRHIDPFGRIHGNYGKILMSPPLNQRRFTRQPEKYMDMSGYTMDFQGRFLDRRGRPVDLLGQHHRHPERFMPPVPYMQQPMPMMPRKFPVVPPFKPMDMSQYTMDFQGRLMDNQGQYVDHMRRFGDHMGPHPPHDQRNLMEFSRYPMDFQDRHFDKYSGPFDRVYKPNTSFERCGSPERGQSPGPETNIHIESRQIIPEKEYTELKSSPGFNRVNSEMSRPSSSGEQXXXXXXXXXXXLSAPPDMESYMFPPIDDEHMQMPFMPPPMFPPIHASMHQPMYSSEKPRMHYPTPEDDPQMQQHMVHFEDEMHMPMQKPFTDDMMMMKMVDMPNAGYQQYQWMDDPFMFPESEYMMDDRQWHSDDGFDYIDDEMPYEYSQIPNMMAVGMHGRPTFRIDEETRDQSHLPPVIGSSDSSEIGRDERCDNRMGYKHMHSMEDREFDSRHGRFETHDDIDYPGFMQDMHQRPMSEFFDHCDSDDGMFDDDYYMDNNMDEDFFPNPRSSMAYPSMGFMNDFQMPFSFDPEYMGMQSGFMPEPEYYPYPYMDDYFGYDKPLVDDKHARYLEDAAEYLKERSRFLRDQHNRFVQELENAGPLKHPQVQAEAEAHANEVKEEVNATEAEAEAVERLARTARRLSNVDHQLAANRHPSLSA |
| 39 | Provided from MBL using Ref^2^ | Esc_cluster_10821p | *Euprymna scolopes* | SQSRTELPSLQVVLSVCPGLPFSSRVAIMNRYMTKHRPMYNQMCRTGRRYRGVMEPMSRMTMDFQGRYMDSQGRMVDPRYYDFSGSSDRYSGRSMFNYGSYMDGGQRYGGFMDNPERYMDMSNYQMDMHGRWMDTQGRYNSPFNYYGYNRHGNYPSYYSYNRNMCNPERMMDMSNYQMDMQGRWMDNYGRHVNPFSHFMYGRNMHYPNFNYYSGRYMDYSDMSNPQMDMQGRYMDSSMSNMYDNFNNYY |
| 40 | Provided from MBL using Ref^2^ | Esc_cluster_1282p | *Euprymna scolopes* | EYMGVSVVNRTPVVSDSLPVPSLPRRFRHLPSVTTMNRYMNRFRNFYGNMYRGRYRGMMEPMSRMTMDFQGRYMDSQGRMVDPRFYDYYGRYNDYDRYYGRSMFNYGWMMDGDRYNRYNRWMDYPERYMDMSGYQMDMYGRWMDMQGRHCNPYSQWMMYNYNRHGYYPNYSYGRHMFYPERWMDMSNYSMDMYGRYMDRWGRYCNPFYQFYNHWNRYGNYPGYYNYYYMYYPERYFDYSNWQMDMQGRWMDMQGRYMDYPYNYYNWY |
| 41 | Provided from MBL using Ref^2^ | Esc_cluster_2045p | *Euprymna scolopes* | MNQIPDRKRFMPHQFYKSEKYPGELEPITMMTMGFQGRYLDSQGRMHDPRVYESYRRYHIGGSNNLNSDPSCSQSYTWWPYPGDYTKPDWYPRPRRQMVFPEKFMDLSSYQMDMKGRWMDMQGRYTNPFNSKNRTRRNHFPLFSPQMYSNNYGNDMSHPEGNKDTPGYPMDSQDHWMYIQGRQTKPSSYNTVGFLGKQYPYKYPPYMHDTWRFISWPERYMDMTGYQMDMRGRWMDTQGRHCNPFNQCGYNKQGSYLGDPYNRNIVYSEKLMDRSNDRMDRQESSDGRYGRRVNPLSRHSCRVCMDMNNPYPGKMDMSNYQMDMQGRWMDTKGRYTNPFPSSGYNKQGPFHGFQHKHNMPYPEKLIDMSNYQMDMQGRWMDTQGRYTNPFSVCGYNRQVPFSGFPYNRNMHYPGKMIDMSNYQMDMQGRWMDTQGRYTNPFSFSSFTRQGPIPGFPYNRNMPYPEKMMDMSNYQMDMQGRWMDTQGRYTNPFSVCGYNRQVPFSGFPYNRNMHYPGKMIDMSNYQMDMQGRWMDTQGRYTNPFSFSSFTRQGPIPGFPYNRNMPYPEKMMDMSNYQMDMQGRWMDTQGRYANPFSLSSFTRQGLFPVFPYNRNMPYPEKMIDMSNYQMDMQGRWMDTQGRYTNPFSFGSFTRQGPIPGFPYNRNMPYPEKMIDMSNYQMDMQGRWMDTQGRYTNPFSFSGYNKQELFPGFSYNRNIAYPENMMDMSNYQMDMDGHWMDTQGRYTNPFGFSSYRRQWPFSGLSYSCNMPYPENMMDMSNYQMDMQGRWMDMYGRYTNPFSFSGFNRRRPFHSFPYNRNMSYPENMMDMSNYQMDMQGRWMDTQGRYRNPFNQYSNYNRYGYFPEYSFDRNTFPMRFNNQIEVPGRFSQPYDHYLSTKDIVYPFYNYRYSHYIYYPDRYMDMSGNHVDMQEPSMDTESRFYPGYQQGRSMNPLNRLLDSHDYNPQTDMEGRWMDYYGEHTHPGFGDQERRTFGDYFRSLPGTYHGDTYRYLGMRNMYDRDQFHYQYGYYQQPPVINSHNDGYYDNNMMGYFYDN |
| 42 | Provided from MBL using Ref^2^ | Esc_cluster_2805p | *Euprymna scolopes* | GPMGMQSGPMGIHGPMGMQGGPMGILGSPMGKQSSPMRMHSGTMRMQSGPMRMRSGPMGMKGGPMGMQSGLMRMRSAPMGMHAFGMNKTSGQQGKPMGRFPRGLRRHMMPMPLSYKRMIRFPERYMDLSGYTMDFKGRFVNQQGKHVNPLHSARMHQGRAMMPAPLPIKRRFMQKPERYMDLSRYTMDFEGRYRDRYGRQIDPMEQFNANIGRYVRNLPPVHRRMVRFPERYMDLSRYTMDFQGRFMDSHGRHIDPFGRIHGNYGKILMSPPLNQRRFTRQPEKYMDMSGYTMDFQGRFLDRRGRPVDLLGQHHRHPERFMPPVPYMQQPMPMMPRKFPVVPPFKPMDMSQYTMDFQGRLMDNQGQYVDHMRRFGDHMGPHPPHDQRNLMEFSRYPMDFQDRHFDKYSGPFDRVYKPNTSFERCGSPERGQSPGPETNIHIESRQIIPEKEYTELKSSPGFNRVNSEMSRPSSSGEQRTEMQYGPQQGNVLSAPPDMESYMFPPIDDEHMQMPFMPPPMFPPIHASMHQPMYSSEKPRMHYPTPEDDPQMQQHMVHFEDEMHMPMQKPFTDDMMMMKMVDMPNAGYQQYQWMDDPFMFPESEYMMDDRQWHSDDGFDYIDDEMPYEYSPIPNMMAVGMHGRPTFRIDEETRDQSHLPPVIGSSDSSEIGRDERCDNRMGYKHMHSMEDREFDSRHGRFETHDDIDYPGFMQDMHQRPMSEFFDHCDSDDGMFDDDYYMDNNMDEDFFPNPRSSMAYPSMGFMNDFQMPFSFDPEYMGMQSGFMPEPEYYPYPYMDDYFGYGFDDDDSEDEREREEEEEESRHRSMKHQTSSSLHRAFHSHGLYHGNMNTRQGVDDVATGIDRPSSLPRIVLSEDKPLVDDKHARYLEDAAEYLKERSRFLRDQHNRFVQELENAGPLKHPQVQAEAEAHANEVKEEVNATEAEAEAVERLARTARRLSNVDHQLAANRHPSLSA |
| 43 | Provided from MBL using Ref^2^ | Esc_cluster_1111p | *Euprymna scolopes* | MFNELDRRYAALQLMLSTDPSHDSRTIANTPKMHHWFATNNRYVSKVMKTKFPITVMIFGVVPSEDTSCRLESQHQRVPGCAEECGGHLVQSFKWPLADSESGSRTQRRPTGPKRPRVAESSEIRDRDPTVMSLSLLLLIHLAYICVWTVNQNQNRKNRAFGGIGIDLEIGFTVSIMNRYFSRHRPMYSHMYGNKYRGMMEPMSRMTMDFQGRYMDSQGRIVDPRFYDFSGSRFNDHDRYYGKSMYGHGSYMDGQRYSGYMDNPERYMDMSNYQMDMYGRWMDMQGRYCSPFTQWSHNRQGNYPGYSYNRNMFHPDRRMDMSNYQMDMQGRWMDMQGRHCSPFNQMGHNRHGNHQWFWHSRYPERWMDMSGYQMDMEGRWMDNYGRYVNPFSNGSYNFGRGMNYPGSYNNYSFGRYMNYPERWMDMSGYQMDMQGPSMDMQGHYMDNFDRNYNDYQMF |
| 44 | Provided from MBL using Ref^2^ | Esc_cluster_1850p | *Euprymna scolopes* | MHQSICGNNAPRLGIVDVHDTHRRRAHFIKLLVEQNTTLRRLYRVSMYRYMNRYQNMLIGHNGKYRSMAEQMSRMSMSPSERMMDPSYYDYYGNGHDDHRYYRGSMYDRDGFWLGNDGHYWYDNWMDNPERYMDMSDYEMDMQGRWMDKHGRYCDPFNQWDCNTYYYYPYHSYGRNMFYPEIYMDMSKYQMDMEGRWMDKKGRYCDPFNDWSYNRYYYYPDYSYFNMLFPERWLDMSSYQMDMEGHWMDLYGRRVNPFSHWMDDGNMYCHQYGFYDDWCMDHPEDWMDMSGYQMDMQGRWMDSKGRYCNPFANFFDCYDMQYHGNNSFFGHLPGNRMSICRYPMDTHGQWMDNQERYDGDY |
| 45 | Provided from MBL using Ref^2^ | Esc_cluster_1112p | *Euprymna scolopes* | YGRWMDMQGRHCSPFNQMGHNRHGNHQWFWHSRYPERWMDMSGYQMDMEGRWMDNYGRYVNPFSNGSYNFGRGMNYPGSYNNYSFGRYMNYPERWMDMSGYQMDMSGRWMDMQGRHCNPYSQWMMYNYNRHGYYPNYSYGRHMFYPERWMDMSNYSMDMYGRYMDRWGRYCNPFYHYYNHWNRSGNNPGYYSYYYMYYPERYFDMSNWQMDMQGRWMDNQGRYCSPYWNNWYGRQMYYPIQNK |
| 46 | Provided from MBL using Ref^2^ | Esc_cluster_9872p | *Euprymna scolopes* | NKYRPMFNNMYSNMYRGRNRGMMEPMSRMTMDFQGRYMDSQGRMVDPRYYDYYGRFNDYDRYYGRSMFNYGWMMDGDRYNRYNRWMDYPERYMDMSGYQMDISGRWMDMQGRHCNPYSQWGYNYNRHGYYPNYSYGRHMFYPERWMDMSGYQMDMQGRYMDRWGRYCNPFSQYMNYYGRYWNYPGYNSYYNSRNMFYPERYFDMSNWQMDMQGRWMDNQGRYCSPYWNNWYGRQMYYPYQNNYFYGRYDYPGMDYSNYQMDMQGRYMDQYGMNDYCY |
| 47 | Provided from MBL using Ref^2^ | Esc_cluster_1281p | *Euprymna scolopes* | MGVSLENRAPFVADCPVLHFSTKVLINMNRYMNRFRNFYGNMYRGRYRGMMEPMSRMTMDFQGRYMDSQGRMVDPRYYDYYGRFNDYDRYYGRSMFNYGWMMDGDRYYRYNRYMDFPERYMDMSGYQMDMYGRWMDMQGRYCNPYSYWMMYNYNRHGYYPNYSYGRHMFYPERWMDMSNYSMDMY |
| 48 | Provided from MBL using Ref^2^ | Esc_cluster_3166p | *Euprymna scolopes* | MNQIPDRKRFMPHQFYKSEKYPGELEPITMMTMGFQGRYLDSQGRMHDPRVYESYRRYHIGGSNNLNSDPSCSQSYTWGPYPGDYTKPDWYPRPRRQMVFPEKFMDLSSYQMDMKGRWMDMQGRYTNPFNSKNRTRRNHFPLFSPQMYSNNYGNDMSHPEGNKDTPGYPMDSQDHWMYIQGRQTKPSSYNTVGFLGKQYPYKYPPYMHDTWRFISWPERYMDMTGYQMDMRGRWMDTQGRHCNPFNQCGYNKQGSYLGDPYNRNIVYSEKLMDRSNDRMDRQESSDGRYGHRVNPLSRHSCRVCMDMNNPYPGKMDMSNYQMDMQGRWMDTKGRYTNPFPSSGYNKQGPFHGFQHKHNMPYPEKLIDMSNYQMDMQAYPENMMDMSNYQMDMDGHWMDTQGRYTNPFGFSSYRRQWPFSGLSYSCNMPYPENMMDMSNYQMDMQGRWMDMYGRYTNPFSFSGFNRRRPFHSFPYNRNMSYPENMMDMSNYQMDMQGRWMDTQGRYRNPFNQYSNYNRYGYFPEYSFDRNTFPMRFNNQIEVPGRFSQPYDHYLSTKDIVYPFYNYRYSHYIYYPDRYMDMSGNHVDMQEPSMDTESRFYPGYQQGRSMNPLNRLLDSHDYNPQTDMEGRWMDYYGEHTHPGFGDQERRTLGDYFRSLPGTYHGDTYRYLGMRNMYDRDQFHYQYGYYQQPPVINSHNDGYYDNNMMGYFYDN |
| 49 | Provided from MBL using Ref^2^ | Esc_cluster_1280p | *Euprymna scolopes* | MDGLPREVHGHVWLPDGHVRTLDGHAGTPLQPVQPMDDVQLQQTRLLSQLLLRPHMFYPERWMDMSNYSMDMYGRYMDRWGRYCNPFYHYYNHWNRSGNNPGYYSYYYMYYPERYFDMSNWQMDMQGRWMDMQGRYCSPYWYNWYGRQMYYPYQNYYWYGRWDYPGMDYSNWQMDMQGRWMDMQGRYMDPWWMNDFSFSLTPRAALLFTSARRSLVRFRLRPNVTTMNRYMNRFRNFYGNMYRNRNRGMMEPMSRMTMDFQGRYMDSQGRMVDPRYYDYYGRYNDYDRYYGRSMFNYGWMMDGDRYNRYNRWMDYPERYMDMSGYQMDMYGRWMDMQGRHCNPYSQWMMYNYNRHGYYPNYSYGRICSTRRDGWTCLTTPWTCTDVTWTGGDVTATRSITITTTGTAPATTPGTIATTTCTTQRDTSTCLTGRWICRDAGWICRDATAAPIGTTGMADRCTTPTRTTIGTADGTIPEWTIPTGRWICRDAGWTCRDDTWTPGG |
| 50 | Provided from MBL using Ref^2^ | Esc_cluster_13228p | *Euprymna scolopes* | DTDKVTIQFSLGERPKIPKNTLVRLPIPSSDTDSGWTATIKETAGSDMTLTVTPPSNCYVGKWRMMGCIVDSDNQTRLSKQKDVCILFNPFNKDDDVYMHDSQEREEYVMNEFGLIFVGTSERIRKRTWNFAQFQGNILDCILYLLDKSEMPYTDRGKPVLVSRRLSAMVNESDDGGVLRGNWSGDYEGGTSPLEWQGSERILEQYWQTKKRVKFGQCWVFSGVLTTACRALGIPCRSVTNFDSAHDTDSSVTIDKFYNENYEYEDMMNEDSVWNFHVWNEAYMTRPDLRDKHYGGWQAIDATPQETSNGIYCCGPCPVRALKNGDIGVNYDASFIMAEVNADSFAWVRQNDGNFIKIQLSSDEIGRYISTKSILKPERNDITHEYKFAEGSAAERVAVLRAVRAGTRADDFHTGIEDVKMEVTLDPTILIGSDFHAKIKIENVSSEVRVVKGNFSVMSVFYTGVLAHSILKHPVKVEIQPGSNELIEIIVKYEEYASKLMDHCGINLTGMLLVDTTNQPLYYNKSYGLEKPDLQIKAPNKGKTGESFDVEVSFKNPLKISLTSCVLRVEGPGLSKGQDLRQKNIGPGETFLTLIQLTPRLPGSRTIHLNFTCDQIAGIDGSHFITIEQS |
| 51 | Provided from MBL using Ref^2^ | Esc_cluster_12890p | *Euprymna scolopes* | RHQVQTIGTRPPSRLGPINSQLLTTDEFKKISEAKDHPSKSDAPMHMPVERQQNFNPPMMAQRAPRYSNPHIGQQGPGMHYDGPRGDVSKQAHKGKKEATEKDRQRPIDNKQRNDGRMESMSKMTMDPQGRYIDSKGKVVDPSTVSDGKHMGSQKRQNTPDDRAPNMPNNDMNFMGPYMDGQDRPYGPYMEPGYMGQMGPMPMRYMDPGHFRMPIDDPERYMDMSRFTMDFEGRYLDPSGQFFEPFGQQGDDPMMDMMTQQMHENNLEEEMTEEELIESGMLVVTDIDYHLPENVRHHHTDQYDLSQNGQLIVRRGQPFMMTIRFNEEYDETKHNLKFIFQIGDNPIPSKKSEVKFGFVDKWHPEEWGAKLVSRQDQFITIYIHAGCDCIVGVWDFLLETMTFGKGSYCFDQFDPMYILFNPWCKGDQVYMNDKDLLCEYILNDHGHIFQGSGSSSVYQKPWNYGQFDDDILDISLYLMRRGFLDCNPQMSNPVRVVRVLAQMINSPENNGVMMRITGADFADGKKPSAWGGSARILQQFIDTKEPVRYGQCWVFAALMTTVCRALGIPCRPVTNFNSNHDSDDNVTVNIYLDENDEGNIYEGKEDNLWSFHVWNDVWMGRPDLPVGYCGWQTIDPTPQEACDGLYCCGPAPVKAIKNGEVNQSYDTNFIFSELNSDRVYWMRNPQSGKWEIVHIEKDALGKYVYTKYPESMPGYSRSGGLMDITNDYKLLCGSEYDRIDIINMSRKKMLMMKAKKSPRQQDDIEFRVTDKENTMVGQKFTVTVSARNTGFDKRSVQTSMYCKVVNHYGAQIGMCREVHINTQFG |
| 52 | Provided from MBL using Ref^2^ | Esc_cluster_10867p | *Euprymna scolopes* | MSTDLNPREHLWGILKQKVEVRKVSNIRQLSDVVIEEWWSIPVATCKALPPTIKMSRQVKTGQGNNTLPKISGNNNSQXXXXTSQTPPKGSHSMSQEGGQSMPQGPQTMPREQPSMSRDPKSTKNKYPMMYEDYGGYSLPPPGSYPMGYSSPPMMSHRGFPMSPDMQSMQGEMYSMYGSPGSNSMSQANNKNRMKTLSKSQRENMFREGPKSGGSYQHSETPKPQTMDRMTMDYEGRFIDRRGRIVDYSRNMDSFDGHRDDKFMFNRDMPRGEQGMCDYGMPAYMYGNMMESPDLMLDMDMYGRYPMFYNQYGDDDEEDYDDSMMDNMINVSHGYMDGRYDTDFAGLNYPDRRTMELVGRYGDSFER |
| 53 | Provided from MBL using Ref^2^ | Esc_cluster_3072p | *Euprymna scolopes* | MPACPDKLLYRKTLDNQGQYLVGIQVLSRGIQVLPRRIQVLSRGIQAFSRGIQVLSREIQVLSRGFQILYKGIQVLSIRLQVPYKRIQVLSRGIQVFFRGITILSKGIQVLSIGILIPYVGIKVQVLSTVCLHVFF*FSRISSLKIHIRTHTGEKPYACKVCEIAFAHNSSLKKHMQTHSRENLNFNAYIGNKNSYGENLNPFGENCNASEENLNASGKNLNAFIGNLKSYGENLNPFIENLKSPGENLNLSGENLNPSGKSLNPSGENLNPSGENLNASRENLNAY |
| 54 | Provided from MBL using Ref^2^ | Esc_cluster_14373p | *Euprymna scolopes* | MSKPQGQRTSHGLKSWPLFSYFNRSRGMPVASPPGPAKDHPSKSDAPMHMPVERQQNFNPPMMAQRAPRYSNPHIGQQGPGMHYDGPRGDVSKQAHKGKKEATEKDRQRPIDNKQSRNDGRMESMSKMTMDPQGRYIDSKGKVVDPSTVSDGKHMGSQKRQNTPDDRAPNMPNNDMNFMGPYMDGQDRPYGPYMEPGYMGQMGPMPMRYMDPGHFRMPIDDPERYMDMSRFTMDFEGRYLDPSGQFFEPFGQQGDDPMMDMMTQQMHENNLEEEMTEEELIESGMLVVTDIDYHLPENVRHHHTDQYDLSQNGQLIVRRGQPFMMTIRFNEEYDETKHNLKFIFQIGDNPIPSKKSEVKFGFVDKWHPEEWGAKLVSRQDQFITIYIHAGCDCIVGVWDFLLETMTFGKGSYCFDQFDPMYILFNPWCKGDQVYMNDKDLLCEYILNDHGHIFQGSGSSSVYQKPWNYGQFDDDILDISLYLMRRGFLDCNPQMSNPVRVARVLAQMINSPENNGVMMRITGADFADGKKPSAWGGSARILQQFIDTKEPVRYGQCWVFAALMTTVCRALGIPCRPVTNFNSNHDSDDNVTVNIYLDENDEGNIYEGKEDNLWSFHVWNDVWMGRPDLPVGYCGWQTIDPTPQEACDGLYCCGPAPVKAIKNGEVNQSYDTNFIFSELNSDRVYWMRNPQSGKWEIVHIEKDALGKYVYTKYPESMPGYSRSGGLMDITNDYKLLCGSEYDRIDIINMSRKKMLMMKAKKSPRQQDDIEFRVTDKENTMVGQKFTVTVSARNTGFDKRSVQTSMYCKVVNHYGAQIGMCREVHINTQFGSKDSKTIMMDVHPEDYLPYMGRNNNSRFSMRISVYCRVKQTNQLFIYSDSFHLDWPYLNVEVPTKFPAGQQVPARISFVNPLPIPLTECELMIQGSGFDRVIEVNLSDVPPHAKMMEEVPCVVRKPGEKQLVATFYCRELSDVVGNAILRVFK |
| 55 | Provided from MBL using Ref^2^ | Esc_cluster_17708p | *Euprymna scolopes* | MKTRDDSVSGIVIRCSFMLCMKQRYVNPGTENQWSGGDNLPYVPFFCVFSAFPDFGHNTYVYCSWCRGLTKNPFIEMKGVRSRLGLSLGKKPKETKTSPTDTKLQTSTMPGRSPLGTGLSSRTNLPLVASNKRGQIPTRDTSRLATNKAPPNPSKEGVGNPLKPNQKTVSKNDYPKSGTNIIQDLLGTQTDSDNSNRIKEELGDNPQTGTGELTGDHVSGLPEEMKDLNTNRDNNDGVQMIHGTTDSDKAEHSLDRHGVNIVKSIQVIHDEDEDDENTDKSSSSSSGKRVDFPTNDDKNTTDNPKRSGSPFIEFLDEDKDTPHPDREEDIVYMYPKGSPPSPEVKRIAAESRSRVDVESTERSNDYFCAEDERLSESPELNLDDRSGEVYFNHKLKKDYSDWSKPYNENTSTYVEDPINYDEFKKNRKFDSDSSEAVRSFKSGYDQKPINYGQLSKNRIDSDFPEDVPSYRPGYDREPFNYGQLNKNRLDSEFSEDVRAYRPGYGPEPINYGQLNKNRRFHSDFPEDVRSYRSGYDRKPINHVQLSKNRFGCDFVDDSDSNRLGNKRILSDSGEFRKNTYGFSTDYEEPYSDYRPNEESRLPNMQDEPTTLNDQLSTRDKYQNEKTSVNPRNLDGEESMRLYEYEFGGHCQSGRNSSQDFEGFTEERENSPERISKIDYIEKKVIQKASARPVQENLQNSQKSQKSDNSQNSSTNFAKNKDANKRETRTSQNNNVKYLHSKGDYDSQTDQNTLRNRNIYSTLKNEDGSSRQLSIFGRWMVTHGNLVDPQNFCPLAYATQPDLYINLSNYEMDAHGRWLDKTGRHVTPFFYCFGYDGEPVYLGDCDYTNGVQDNYMNDNDYNDDFVNNPERYIDLSDYQMNMDKQWVNSLGRDIDPLSDWYSDGSEGDEHNEYSWMYDRPQFLEDSGLPNPMDGRYSNFYQPDTPYFNDTNDRFPMGEAFSETCEYWDFAEQYGCMENGIESMQHPWQAAGDFSRGYTGANEEYFVEEPDSVPYQQRSSKDSDK |
| 56 | Provided from MBL using Ref^2^ | Esc_cluster_18323p | *Euprymna scolopes* | MESAPRLARFRSAIDEASANQVNAVILKDRRVTICQKAQKFKINIGFENHPLLEMNTFMDTMHCDGMGMPQSKFGDFSHNCMRSFPKSQRELMRRDLVAKPGKNRRFGDFMEPMSRMTMDFCGRMIDSQGRIVDPSRFLFMEEHYMDNDRFPYFYDMMRSPRSMYSGMYGFGSGDHSFNRGMFNDDMYRDMYHGGMNPFMYNRSMMGRMYSPGRFMDDSFSMYYRPRMGDHFMYSQSQFNDQEGGQGMFGRMSDNFETSPGRPTEEQSIARRLSESHNLHRRLSESQARIEAANNQRKASRALIFPEETTNMESA |
| 57 | Provided from MBL using Ref^2^ | Esc_cluster_18324p | *Euprymna scolopes* | MYYDGTCVPYQNFGYNYTRGFPKLHRDTMRRDLMVTSGKNRTFEDFMELMSGMTMDFGGRMIDSQGRIVDPSFFDEYYIDYDRFPYFHDTMRSPRFMYSGMYGFGSGDHSFYRSMYNDDMYRDMYHGGMNPFMQNRSMMGRMYSPSRFMDDPFSMRYRPRMSYHLMDSQNQFNDQEVGQGMFGRMSHNIGRSIGRPLEEQKLILVTATNITRRNPKL |
| 58 | “Reflectin, Reflectin 8” from Uniprot | I0JGV7 | *Sepia officinalis* | MNRFMNRYRPMFNNMHNNMYNNMYRGRYRGMMEPMSRMTMDFQGRYMDSQGRMVDPRYYDYYGRWNDYDRYYGKSMFNYGWMMDGDRYNNYYRWMDFPERYMDMSGYQMDMYGRWMDMQGRHCNPFNQWGHNRYGQSFNYNYGRNMFYPERWMDMSNYSMDMQGRYMDRWGRHCNPFSQNMNWYGRYWNYPGYNNYYYNRHMYYPERYFDMSNWQMDMQGRWMDMQGRHNNPYWYNWYGRQMYYPYQNNWYGRWDYPGMDYSNWQMDMQGRWMDMQGRYMDPWMSDYSYNN |
| 59 | “Reflectin” from Uniprot | I7LGW2 | *Sepia officinalis* | MNRYMNRFRNWYGNNYRGRYRGMMEPMSRMTMDFQGRYMDSQGRMVDPRYYDYYGRWNDYDRYYGRPMFNYSWMMDGDRYNRYYRWMDFPERYMDMSGYQMDMYGRWMDMQGRHCNPFRQWGYQRHGYYPGYHYGSNMFYPERWMDMSNYSMDMQGRYMDRWGRHCNPFSHYYNHWNRYWNHPGYYNYYYMYYPERYYDMSNWQMDMQGRWMDMQGRHCNPYWYNWHGRHMYYPYQNYYWYGRWDYPGMDYSNWQMDFQGRWMDNQGRYMDPWWWNDYYSYYY |
| 60 | “Reflectin” from Uniprot | I7KQN6 | *Sepia officinalis* | MNRYTMRNRPMYGNMYRTGKKYRGVMEPMSRMTMDFQGRYMDSQGRMVDPRHNDYYGRWNDYDRYYGRSMFNYGPHMDGHQHGGWMDFPERWMDMSNYQMDMQGRWMDMQGRHCQPFNQWGYNRHGNYPSSYYGRNMFYPERWMDMSNWQMDTQGRWMDMQGRYGSPFNQWGYNRHGYYPGSSYGRNMYHPERWMDMSNYQMDMQGRWMDMHGRHVNPFSHSMHGRNWSYPYYNYYSSRHMDYPERNMDMSNWQMDMQGRWMDMQGRHMDPSWSNMHDNHNYWF |
| 61 | “Reflectin” from Uniprot | I7LE98 | *Se Sepia phia officinalis* | MNRSMNRWRPMFNNMHNNYYGRSMFNYNWMMDGDRYNRYYRWMDFPEWYMDMSGYQMDMYGRWMDMHGRHCNPFRQWGYQRHGYYPGFHYGSNMFYPERWMDMSNYSMDMQGRYMDRWGRHCNPFSHYYNHWNRYWNHPGYYNYYYMYYPERCYDMSNWQMDMQGRWMDMQGRHSNPYWYNWHGRHMYYPYQNYYWYGRWDHHGMDYSNWQMDFQGRWMDNHGRHMDPWWNEHYFNHYY |
| 62 | “Reflectin” from Uniprot | I7J3R4 | *Sepia officinalis* | MNRFMNRWRPMFNNMRNNMYRGRYRGMMEPMSRMTMDFQGRYMDSQGRMVDPRYYDYYGRWNDYDRYYGRSMFNYSWMMDGDRYNRYYRWMDFPERYMDMSGYQMDMYGRWMDMHGRHCNPFRQWGYQRHGYYPGFHYGSNMFYPERWMDMSHYSMDMQGRYMDRWGRHCNPFSHYYNHWNRYWNHPGYYNNHYNRHMYYPERYFDMSNWQMDMQGRWMDMQGRHSDPYWYNWHGRHMYYPYQNYYWYGRWDHHGMDYSNWQMDFQGRWMDNHGRHMDPWWNEHYFNHYY |
| 63 | “Reflectin” from Uniprot | I7K2G8 | *Sepia officinalis* | MNRSMNRWRPMFNNMHNNYYGRSMFNYDWMMDGDRYNRYYRWMDFPERYMDMSGYQMDMYGRWMDMQGQHYNPFRQWGYQRHGYYPGYHYGSNMFYPERWMDMSNYSMDMQGRYMDRWGRHCNPFSHYYNHWNRYWNHPGYYNNHYNRHMYYPERYFDMSNWQMDMQGRWMDMQGRHCNPYWYNWHGRHMYYPYQNYYWYGRWDYPGMDYSNWQMDFQGRWMDNHGRHMNPWWNEHYFNHYY |
| 64 | “Reflectin 10” from Uniprot | I0JGV9 | *Sepia officinalis* | MNRFMNRWRPMFNNMRNNMYRGRYRGMMEPMSRMTMDFQGRYMDSQGRMVDPRYYDYYGRWNDYDRYYGRSMFNYSWMMDGDRYNRYYRWMDFPERYMDMSGYQMDMYGRWMDMHGRHCNPFRQWGYQRHGYYPGFHYGSNMFYPERWMDMSHYSMDMQGRYMDRWGRHCNPFSHYYNHWNRYWNHPGYYNNHYNRHMYYPERYFDMSNWQMDMQGRWMDMQGRHSNPYWYNWHGRHMYYPYQNYYWYGRWDHHGMDYSNWQMDFQGRWMDNHGRHMDPWWNEHYFNHYY |
| 65 | “Reflectin 6” from Uniprot | I0JGV5 | *Sepia officinalis* | MNRSMNRWRPMFNNMHNNYYGRSMFNYNWMMDGDRYNRYYRWMDFPEWYMDMSGYQMDMYGRWMDMHGRHCNPFRQWGYQRHGYYPGFHYGSNMFYPERWMDMSNYSMDMQGRYMDRWGRHCNPFSHYYNHWNRYWNHPGYYNYYYMYYPERYYDMSNWQMDMQGRWMDMQGRHSNPYWYNWHGRHMYYPYQNYYWYGRWDHHGMDYSNWQMDFQGRWMDNHGRHMDPWWNEHYFNHYY |
| 66 | “Reflectin 7” from Uniprot | I0JGV6 | *Sepia officinalis* | MNRFMNRYRPMFNNMHNNYYGRSMFNYNWMMDGDRYNRYYRWMDFPERYMDMSGYQMDMYGRWMDMQGRHCNPFRQWGYQRHGYYPGFHYGSNMFYPERWMDMSNYSMDMQGRYMDRWGRHCNPFSYYYNHWNRYWNHPGYYNYYYMYYPERYYDMSNWQMDMQGRWMDMQGRHSNPYWYNWHGRHMYYPYQNYYWYGRWDYPGMDYSNWQMDFQGRWMDNHGRHMNPWWNEQYFNHYY |
| 67 | “Reflectin 1” from Uniprot | I0JGV0 | *Sepia officinalis* | MNRYMMRNRPMYGNMYRTGKKYRGVMEPMSRMTMDFQGRYMDSQGRMVDPRHNDYYGRWNDYDRYYGRSMFNYGPHMDGHQHGGWMDFPERWMDMSNYQMDMQGRWMDMQGRHCQPFNQWGYNRHGNYPSSYYGRNMFYPERWMDMSNWQMDTQGRWMDMQGRYGSPFNQWGYNRHGYYPGSSYGRNMYHPERWMDMSNYQMDMQGRWMDMHGRHVNPFSHSMHGRNWSYPYYNYYSSRHMDYPERNMDMSNWQMDMQGRWMDMQGRHMDPSWSNMHDNHNYWF |
| 68 | “Reflectin 5” from Uniprot | I0JGV4 | *Sepia officinalis* | MNRFMNRYRPMFNNMHNNYYGRSMFNYDWMMDGDRYNRYNRWMDFPERYMDMSGYQMDMYGRWMDMQRHHCNPFRQWGYQRHGYYPGFHYGSNMFYPERWMDMSNYSMDMQGRYMDRWGRHCNPFSHYYNHWNRYWNHPGYYNNHYNRHMYYPERYFDMSNWQMDMQGRWMDMQGRHSNPYWYNWHGRHMYYPYQNYYWYGRWDHHGMDYSNWQMDFQGRWMDNHGHHMNPWWNEQYFNHYY |
| 69 | “Reflectin 2” from Uniprot | I0JGV1 | *Sepia officinalis* | MNRFMNRYRPMFNNMHNNYYGRSMFNYDWMMDGDRYNRYYRWMDFPERYMDMSGYQMDMYGRWMDMQGHHCNPFRQWGYQRHGYYPGFHYGSNMFYPERWMDMSNYSMDMQGRYMDRWGRHCNPFSHYYNHWNRYWNHPGYYNYHYNRHMYYPERYFDMSNWQMDMQGRWMDMQGRHSNPYWYNWHGRHMYYPYQNYYWYGRWDHHGMDYSNWQMDFQGRWMDNHGRHMNPWWNEQYFNHYY |
| 70 | “Reflectin 3” from Uniprot | I0JGV2 | *Sepia officinalis* | MNRFMNRYRPMFNNMHNNYYGRSMFNYDWMMDGDRYNRYYRWMDFPERYMDMSGYQMDMYGRWMDMQGRHCNPFRQWGYQRHGYYPGFHYGSNMFYPERWMDMSNYSMDMQGRYMDRWGRHCNPFSQYYNHWNRYWNHPGYYNNHYNRHMYYPERYFDMSNWQMDMQGRWMDMQGRHSNPYWYNWHGRHMYYPYQNYYWYGRWDHHGMDYSNWQMDFQGRWMDNHGRHMDPWWNEQYFNHYY |
| 71 | “Reflectin 4” from Uniprot | I0JGV3 | *Sepia officinalis* | MNRSMNRWRPMFNNMHNNYYGRSMFNYDWMMDGDRYNRYYRWMDFPERYMDMSGYQMDMYGRWMDMQGQHYNPFRQWGYQRHGYYPGYHYGSNMFYPERWMDMSNYSMDMQGRYMDRWGRHCNPFSHYYNHWNRYWNHPGYYNNHYNRHMYYPERYFDMSNWQMDMQGRWMDMQGRHCNPYWYNWHGRHMYYPYQNYYWYGRWDYPGMDYSNWQMDFQGRWMDNHGRHMDPWWNEQYFNHYY |
| 72 | “Reflectin 9” from Uniprot | IOJGV8 | *Sepia officinalis* | MNRYMNRFRNWYGNNYRGRYRGMMEPMSRMTMDFQGRYMDSQGRMVDPRYYDYYGRWNDYDRYYGRSMFNYSWMMDGDRYNRYYRWMDFPERYMDMSGYQMDMYGRWMDMQGRHCNPFRQWWYHRHGYYPGFHYGSNMFYPERWMDMSNYSMDMQGRYMDRWGRHCNPFSHYYNHWNRYWNHPGYYNYYYMYYPERYYDMSNWQMDMQGRWMDMQGRHSNPYWYNWHGRHMYYPYQNYYWYGRWDYPGMDYSNWQMDFQGRWMDNQGRYMDPWWWNDYYYNYYY |
| 73 | “Reflectin 11” from Uniprot | IOJGW0 | *Sepia officinalis* | MNRFMNRWRPMFNNMHNNMYRGRYRGMMEPMSRMTMDFQGRYMDSQGRMVDPRYYDYYGRWNDYDRYYGRPMFNYSWMMDGDRYNRNYRWMDFPERYMDMSGYQMDMYGRWMDMQGHHCNPFRQWGYQRHGYYPGFHYGSNMFYPERWMDMSNYSMDMQGRYMDRWGRHCNPFSHYYNHWNRYWNHPGYYNNHYNRHMYYPERYFDMSNWQMDMQGRWMDMQGRHSNPYWYNWHGRHMYYPYQNYYWYGRWDHPGMDYSNWQMDFQGRWMDNHGRHMNPWWNEHYFNHYY |
| 74 | “Reflectin_like” from Uniprot | I0JGW1 | *Sepia officinalis* | MSTFMDPMFYEGLGMPPPNFGDFNHNCMRSFHKSQRDMMRRDIMAKSSKNKRCGDLMEPMSRMIMDFNGGLIDSRRRITDSDHYFTIDGNYGDNEKPLTSDGLLRNRYDMYAFAPADKCHNRTRGLYGDSMYRDKHQDGMYSSGYMQGRSMQNHRMMGSFQGGMQTQSRYMDDPYYVNYNSGTYDTPVDMNSYYFDQEGRHRMYSRFSEGQITPGRQEGTYSARRESRSGQRRLSDSHSFQRPIDTRSNRRTSHGMLYSERNNIDFA |
| 75 | “Reflectin, Reflectin 12” from Uniprot | A0A812C5C9 | *Sepia pharaonis* | MNRFMNRWRPMHNNMYRGRYRGMMEPMSRMTMDFQGRYMDSQGRMVDPRYYDYYGRWNDYDRYYGRSMFNYRWMMDGDRYNRYNRWMDFPERYMDMSGYQMDMYGRWMDMHGRHCNPFNQWWNQRHGYYPGFHYGSNMFYPERWMDMSNYSMDMQGRYMDRWGRHCNPFSQYMNWYGRYWNYPGYNNYSYNRHMYYPERYYDMSNWQMDMQGRWMDMQGRHSNPHWYNWYGRQMYYPYQNNWYGRWDYPGMDYSNWQMDMQGRWMDNHGRHMDPWMNDYSYNNW |
| 76 | “Reflectin, reflectin 1” from Uniprot | A0A812CAZ1 | *Sepia pharaonis* | MNRYMMRHRPMYSNMYRTGKKYRGMMEPMSRMTMDFQGRYMDSQGRMVDPRHYDYYGRWNDYDRYYGRSMFNYGPHMDGQRHGGWMDFPERWMDMSNYQMDMQGRWMDMQGRYCHPFNQWGYNRHGNYPGYSYGRNMFYPERWMDMSNWQMDTHGRWMDMQGRYGSPFSQWGYNRHGYYPGYSYGRNMFYPERSMDMSNYQMDMQGRWMDMQGRHVNPFSHSMYGRNWSSPYYNYYSSRHMDYPERFMDMSNWQMDTQGRWMDMQGRHMDPSWSNMQDNYNNWF |
| 77 | “Reflectin, reflectin 14” from Uniprot | A0A812C8L7 | *Sepia pharaonis* | MNRFMNRWRPMFNNMHNNYYGRSMFNYDWMMDGDRYNRYYRWMDFPEWYMDMSGYQMDMYGRWMDMQGRHCNPFRNWWHQRHGHYPGFHYGGNMFYPERWMDMSNYSMDMQGRYMDRWGRHCNPFSQFYNHWNRYWNQPGYYNYHYNWHMYYPESYFDMSNWQMDMQGRWMDRQGRYSNPYWYNWQGRHMYYPYQNYYWYGRWNYPGMDYSNWQMDFQGRWMDNHGRHMDPWWNEHYFNHYY |
| 78 | “Reflectin, reflectin 9” from Uniprot | A0A812BXX5 | *Sepia pharaonis* | MNRYMNRFRNWYGNNYRGRYRGMMEPMSRMTMDFHGRYMDSQGRMVDPRYYDYYGRWYDYDRYYGRPMFNYSWMMDGDRYNRYYRWMDFPERYMDMSGYQMDMYGRWMDMQGRHCNPFRHWWHQRHGYYPGFHYGSNMFYPERWMDMSNYSMDMQGRYMDRWGRHCNPFSHFYNYWNRYWNHPGYYNYYYMYYPERYYDMSNWQMDMQGRWMDMQGRHNNPYWYNWQGRHMYYPYQNYYWYGRWDYPGMDYSNWQMDFQGRWMDMQGRYMDPWWWNDYYYNYYYY |
| 79 | “Reflectin, reflectin 13” from Uniprot | A0A812C2Y5 | *Sepia pharaonis* | MNRFMNRWRPMHNNMYRGRYRGMMEPMSRMTMDFQGRYMDSQGRMVDPRYYDYYGRWNDYDRYYGRPMFNYRWMMDGDRYNRYNRWMDFPERYMDMSGYQMDMYGRWMDMHGRHCNPFNQWWHQRHGYYPGFHYGSNMFYPERWMDMSNYSMDMQGRYMDRWGRHCNPFSHFYNHWNRYWNQPGYYNSHYNWHMYYPERYYDMSNWQMDMQGRWMDMQGRHSNPYWYNWHGRHMYYPYQNYYWYGRWNYPGMDYSNWQMDFQGRWMDNHGRHMDPWWNEHYFNHYY |
| 80 | “Reflectin, reflectin 14” from Uniprot | A0A812BXY5 | *Sepia pharaonis* | MNRFMNRWRPMFNKMHNNYYGRSMFNYDWMMDGDRYNRYYRWMDFPEWYMDMSGYQMDMYGRWMDMQGRHCNPFRNWWHQRHGHYPGFHYGSNMFYPERWMDMSNYSMDMQGRYMDRWGRHCNPYSQFYNHWNRYWNQPGYYNSHYNWHMYYPESYFDMSNWQMDMQGRWMDRQGRYSNPYWYNWQGRHMYYPYQNYYWYGRWNYPGMDYSNWQMDFQGRWMDNHGRYMDPWWNEHHFNHYY |
| 81 | “Reflectin, reflectin 9” from Uniprot | A0A812C8S0 | *Sepia pharaonis* | MNRFMNRFRNWYGNSYRGRYRGMMEPMSRMTMDFHGRYMDSQGRMVDPRYYDYYGRWNDYDRYYGRSMFNYRWMMDGDRYNRYNRWMDFPERYMDMSGYQMDMYGRWMDMHGRHCNPFNQWWHQRHGYYPGFHYGSNMFYPERWMDMSNYYMDMQGRYMDRWGRHCNPFSHFYNHWNRYWNQPGYYNYYYMYYPERYYDMSNWQMDMQGRWMDMQGRHSNPYWYNWHGRHMYYPYQNYYWYGRWDYPGMDYSNWQMDFQGRWMDNYGRYMDFPFYNWY |
| 82 | “Reflectin, reflectin 14” from Uniprot | A0A812C3T1 | *Sepia pharaonis* | MNRFMNRWRPMFNNMRNNYYGRSMFNYDWMMDGDRYNRYYRWMDFPEWYMDMSGYQMDMYGRWMDMQGRHCNPFRHWWHQRHGHYPGFHYGGNMFYPERWMDMSNYSMDMQGRYMNRWGRHCNPFSHFYNHWNSYWNHPGYYNYHYNWHMYYPESYFDMSNWQMDMQGRWMDRQGRYSNPYWYNWQGRHMYYPYQNYYWYGRWNYPGMDYSNWQMDFQGRWMDNHGRYMDPWWNEHHFNHYY |
| 83 | “Reflectin, Reflectin 14” from Uniprot | A0A812C5Z0 | *Sepia pharaonis* | MNRFMNRWRPMFNNMHNNYYGRSMFNYDWMMDGDRYNRYYRWMDFPEWYMDMSGYQMDMYGRWMDMQGRHCNPFRHWWHQRHGHYPGFHYGGNRFFPERWMDMSNYSMDMQGRYMNRWGRHCNPFSHFYNHWNRYWNHPGYYNYHYNWHMYYPESYFDMSNWQMDMQGRWMDRHGRHSNPYWYNWHGRHMYYPYQNYYWYGRWNYPGRDYSNWQMDFQGRWMDNHGRYMNPWWNEYYFNHYY |
| 84 | Provided from MBL using Ref^3^ | Aargo020156_Reflectin | *Argonauta argo* | MNRLMNRFRRQFGRKYKGFMEPMNMMSMDFQGRYMDSYGRMVNPRYYEYYGRYSDQDRYYGRSMYNYYGFYDNDRYNRQGYFMDFPERFMDMSGYQMDMYGRWMDMQGQHSSPYGHMFYSSRHGYYPGYQYGRNYGYPERFMDMSYYQMDIYGRYMDRFGRFCNPYYNYYRRYMYYPFMNFYYSYYPERFMDMSNYQMDMYGRWMDMYGRYSSPYYSYYGRYYHNYPYYSYSWGNRHYSYPERYFDMTNYQMDFEGRWMDMYSRHCTPFYNYYGRYHHNYPSYNYSWFQKSFSYPDRYYDMDYESHSMSMYGYNYPYNSYSRHYYFNYSPYYNYFGGNRYYNNYDRFFDMSNYQMDFSGQWMDMNGGYLSHFDRWNEYYFY |
| 85 | Provided from MBL using Ref^3^ | Aargo020155_Reflectin | *Argonauta argo* | MNKYMNRPQNQFGGKYRGFMEPMNMMSMDFQGRYMDSYGRMVDPRNYGYYGRYSDQDRYYGRSMYNYYGLYDNDRYNRQGYFMDFPERFMDMSGYQMDMYGRWMDMQGQHSSPYGHMFNSSRHGYYPGYQYGRNYGYPERFMDMSYYQMDIYGRYMDRFGRFCNPYYNYYRNYMYYTFMNFYYSYYPERFMDMSNYQMDMYGHWMDMYGRYSSPYYSYYGRYYHNYPYYSYSWGNRHYSYPEKYFDMANYQMDFEGRWMDMYSRHCTPFYNYYGRYHHNYPSSDNMWGQRSFNYPDSRFEMDYQPFNQMYNSYGNHIDNYRYKGNTSGPYYFGYYSYYPYQSQRNNPDMNVNQLEIQYNSMDLNNHNRDQINVVKNSDYHSDFEKQWM |
| 86 | Provided from MBL using Ref^3^ | Aargo020153_Reflectin | *Argonauta argo* | MNKFFNMSRRNNFSRRYRGIMEPMSKMTMDFQGRYMDSYGRMVDPRNYGYYGRYSDQDRYYGRSMYNYYGFYDNDRYNRQGYFMDFPERFMDMSGYQMDMYGRWMDMQGQHTSPYWHMFNSSRHGYYPGYQYGRNYGYPERFMDMSYYQMDIYGRYMDRFGRFCNPYYNYYRRYMYYPFMNFYYSYYPERFMDMSNYQMDMYGRWMDMYGRYSSPYYSYYGRYYHNYPYYSYSWGNRHYSYPERHFDMANYQMDFEGRWMDMYSRHCTPFYNYYGRYHHNY |
| 87 | Provided from MBL using Ref^3^ | Aargo019690 | *Argonauta argo* | MVTLCCTSGLTGLYQWFNYGLIIVNQSSNCGLIIVYWCVGVGVVASVGVVVGGVSVGGVGVGVVNVGGGGVGGGIVGVGVVVVSVGVVVVNVGVGGGIVGVVVVVVSVGVGCVINVGVVVNVGGDGVVVSVVVGVGVGVVVVVVKMLYDSQNGENNPVYLRLRRPVDSGSHHMEDRRSNENILASSAGFPYDNATILTSSARNIASDGSRLLPSYLDRQNLLNRGKFSYPESMIQRDVSQLNNSQQQIFGMKQDTFDRQARKSLYFQGGGGGGDDNTDDTLLVTYLSRKGGSPFSPQSFHEIAANCPEIKVDDQMNTFNKRRSHIEPLENPLVLSGNKKTLVDNSLALVQFPSQDGQVSIRQPTSLPTDRLLHSQNNFPQQSNFSYFPRKQFFNQFASPYFDRGNQFARFMPNRPPFSAGYPSYMNNPFSQNRPSFRMGPRCEYMPYMQSNQCPCQECCSLSGRQKPSGNFDWDCMNDPDYDQYPWNQNDPTYMDLPNYPGKNSQNNFGFGPPNCCRDTSYFQDLPSNMYNIPPSTLTNIDEIDPTSDAYLEKIIPNFNILPPKSNLKIGPMAPELAENYVKNYLFVPSNQTRIISPENSPSEVTHQTSRSTMLSRLATLLERSIDMSSYSMNFDGNYIDNRGLLVDPFQTYITDLEQGHSKPQGGGGRGRRSGRERTAPETMSTQPSVVILRNSRSDVMTLDEGDILKNTAKRVEENEDETKDKMNDVSFLTNLTRRMDLHDKPQQEHEDMAGPSQQNIFQSHPALIAEAMRRENNVRDDYCVVPPYYPAPGARERYFQPPAAAEGPIDMMPLWEQERIAREREFHRYKSQFNPGNRMFLNTTTPLIKSEIYDDRMDARNMTTAPQQGVGRDREGADETSYRTMADYLQSPAAKVNPRLLNKLNYQNKTQSSKDVPFVTEASDDSDTEREKAQQEEVIGADTFARLKNRFETERRNKLNGEDNTGVRRKHRYFRGLEGGKLLQGPGAGNCPY |
| 88 | Provided from MBL using Ref^3^ | Aargo020154 Note=XP_029640887.1(TBC1 domain family member 5 homolog A-like) | *Argonauta argo* | MEPMSRMTMDFQGRYMDSSGKLVDPRYNDYNGRYSNYDRFGNRPMYGGGFPDNDKFQKYGRFMNFPEKHMDMSGYQMDMRGRYMDKYGRYCNPYSRRHMNYPSNNYDNQQMYNGEKFMDLSNYQMDMHGRWMDSNGRYSAPFNNFGGRYFHNYPQPPFMGPRGLNHPDRFFDMSNYQMDFDGKWMDMHGRHSHPFNDNHFGRQHYMQPHHNYPQGHRHFNNPERYIDMGNYQMDFDGKWMDMQGRHFHPFGGNNSHHARYHHSYPYNSFNHGHRYFNNPEKYFDMTNYQMDFDGKWMDMQGRHSHPYAGNNSHNVRHHHSYPYNSFNQGHRYFNNPERYFDMTNYQMDFDGKLMDMQGRHFHPFSGNNNYHGRHFHNQPSNSFGHGHRYTSNVDRNFDMGNYQMDFEGRWMDMHGRHSHPYQGHNFQSRHQHNSPHHFSWGHKYSHYPEKFFDMGNYQMDFDGQWMDMEDRHCLPFSGNLNQSNRYSQHYNQFPNHNFHWGQRNQDHPERYFDMSGYQMDFDGRWMDVNTSSHHDNFFQM |
| 89 | Provided from MBL using Ref^3^ | >Aargo020157 Note=XP_029640363.1(hemocyte protein-glutamine gamma-glutamyltransferase-like isoform X1) | *Argonauta argo* | MDKRHQGNKDRLKHWPFLNYFNRSRNMPPVCPPSDTNNKEKTDSMGRMPRGNARHFGPPFMGPPMPGECFEGLPPPPPCVMNSLSRKKRIPEKERMEDMDGHSKFEDSTEKRPKSPTMSLGQKSECRDKEPRDPIQEDLRDRNNPCDQKPEIPSQHRDSLVFDTESKCSDLEKTNGFYPYPDFDSHGMMPPFGMMPMGYGPPPDGMRHPMYMNYPEKPIDMSRLTMDFEGKYMDDNGQFIDPLEMQDGNQMDIPRENMNFRPQYMEQYPGDMDDCMGISPMEMKRYNHRRRDGYDEMSPQQLTENGLLIVTNVDYHLDESLRLHHTEQYDIGQNGQLIVRRGQPFMLSIEFQQEFDLERHKLRFIFQIGEHPLPSKKSDVRFSLTDKWHPEEWGAKLVGRRGPMITVCIHPASDCIVGVWDFMIKTIVNGKGSYVFDNFDPIYILFNPWCQNDQVYMEDKELLYEYVLNDHGYIFQGSGCSSVYQKPWNYGQFDEDILDIALYLVRQGFLEHCPQMSSPVRIARVLVHLINSPENTGVMMRFDTDECQNGKRPTAWGGSRRILQQFIDTKEPVRYGQCWVFSGLLTTVCRALGIPCRTVTNFNSNHDSDNNLSVNVYLGENEKGNIFEQKDGNVWNFHVWNDVWMSRSDLPVGYCGWQAVDPTPQEASDGVYCCGPAPLKAIKNGEVNQNYDASFIFSELNADRLYWMKNLRSGKWEIVHIDRNAMGKFIYTKYPNCKPGHNRSGGLMDVTYDYKMINASEYDRIDVINVHRKKLMLQKAKKPQHYQEDMEFRLTERESVMIGQDFNVTLQCRNIGQEQRYVNCSMYCKIVDHFGDKIAICRDIHVNNFFDPKSNKMVVMNVSPDDYMPFLGRDNNSRLSMRIMIYCKVKQTEQMFIFNDSYQLDWPYLNIEVPPKILLRDEVRARISFINPLSISLTNCEFMVEGNGFERIYETRVNDIPPHGNFMEEILFVGRKIGEKQLVATFYCQELCDVVGNAILQVYKN |
| 90 | Provided from MBL using Ref^4^ | Ocbimv22030998m_Scaffold57339_45977-54246 | *Octopus bimaculoides* | MNRSRNMFRNSSRKHRGVMEPMTRMTMDFQGRYLDSSGRLVEPRCNDYYGRNSNYDRYRPMQNTGIYDNDKFQKYGRFMHFPERQMDMSGYQMDMRGRYMDKYGRHCNPYSRRHMNYPNNNYDNYHMYNPEKLMDMSNFQMDMHGRWMDSNGRYSSPFSNYGSRHHQNYPHFNYNWGQRGFNYPDRFFDMSNYQMDLDGKWMDTYGRHCHPFYDNSNYYGKQYNYNMYPHYNYNWGQKYYHYPERYFDMSNYQMDFDGRWMDMFGRSHSPFNGYNNNQGRQHHGQPHNSFSYGQRYQDRNFDIGNYQMDFDGRWMDMYGRYSHPFYGYNNFQSRYQHNLPQNFNWGQRSFHNPERLFDMGNYQMDFDGHWMDMDDRHCQPFTGNYNHSNRYQQNCNHSPSQNFNWSQRYQDNPEKFFDMSGYQMEFDGRWMDSNNYNSDNFW |
| 91 | Provided from MBL using Ref^4^ | Ocbimv22031000m_Scaffold57339_101048-107169 | *Octopus bimaculoides* | MNRLMNKFRHHFGRKYRGIMEPMSVMSMDFQGRYMDSYGRMVDPRFYEFYGRYSDNDRYYGKSMYNYYGFYDNDRFHRYGNFMDFPERFMDMSSYQMDMYGRWMDMHGHHSSPYWYMFNSSRHGHYPGYRYGRNWFYPERFMDMSHYQMDMYGRYMDRYGRQCNPYYNYYRRYMYYPYMNFYYMHYPERFMDMSGYQMDMYGRWMDMYGRHSTPFYTNYGRYYHNYPYYNYSWGQRYYNYPERYFDMTNYQMDFDGRWMDMYSRHCTPFYSYHGRYHHYYPYHSYSWGQRFYNNPERWYDMDYEFHSMSPYNYHSRYHYFNYSPYFSGGHRWFDMSNYQMDFGGQWMDMNGRYMNHFDHWNEYFF |
| 92 | Provided from MBL using Ref^4^ | Ocbimv22030997m_Scaffold57339_15001-19853 | *Octopus bimaculoides* | MNRYMNRRNNFSRRYRGIMEPMSRMTMDFQGRYMDSYGRMVDPRFYGFYGRYSDNDRYYGRSMYNYYGFYDNDRFHRYGNFMDFPERFMDMSGYQMDMSGRWMDMHGHYSSPHWHMFNSSRQGYYPGYHYGRNWFYPERFMDMSHYQMDMYGRYMDRNGRHCNPYYNYYRRYMYYPYMNFYHMYYPERFMDMSGYQMDMYGRWMDMSGRHSSPFYSYHSRFHHNYPYYNYSWGQRYYNYPERYFDMGNYQMDFDGRWMDMYSRHCTPFYNYHGKFHHNYPYYNYSWGQRYYNYPERYFDMGNYQMDFDGRWMDNYGRYSSPFNSYHGRFHHNYPYYNYSWGQRYYNYPERYFDMGNYQMDFDGRWMDNYGRYSSPFYNYHGRYHNYPYYNYSWGQRYYNYPEGNYQMDFDGRWMDNYGRYFHGYNYHNRHYYNSYPNSYNYNWGQRYYDYPERNFDMFNYQMDFDSRWMDGQNFHYYGDNYNY |
| 93 | Provided from MBL using Ref^4^ | Ocbimv22030999m_Scaffold57339_81227-85638 | *Octopus bimaculoides* | MNRFMNRFRPQFNRKYRGFMEPMNMMSMVFQGRYMDSYGKMVDPKLYEFYGKYSDNDRYYGKSMYNYYGFYDNDRFHRNGNFMDFPERFMDMSGYQMDMNGKWMDTQGQNSHPYWNMFSSSRQGCYPGYSYGRNWFFPERFMDMSHYQMDMNGRYMDKSGRHCNPYYSYYRRYMSHPQMNFNQMHYPERFMDMSSYQMDIGGRYMDKWGCHINPFSTYYFGKQSYFPHNYWSQRKYMDMSSYQMDMQYNSMDMNSRNCDQLHYFRNFDMWNNQMDFDGHWMNMNNQSYHPSSFIRNQVYYNPYHFYTWMSRYYNHPEKFYDTSNYQVEFGGKWPSQEYECITQE |
| 94 | Provided from MBL using Ref^4^ | Ocbimv22030996m_Scaffold57339_1379-1806 | *Octopus bimaculoides* | MFYDMSHYQMDFDGNWMDMYNHYSHPFFGYDHYHRGSHYYNHYPYHNYSWGHRFYDYPERFFDMSHYEMDFNGRWMNMHRF |
| 95 | Provided from MBL using Ref^4^ | Ocbimv22014010m_Scaffold212420_23002-26887 | *Octopus bimaculoides* | MYGHIYNGRMEPMHMMSMDFHGRYMDSYGRMVDPRSYGFYGRYHDQDSYYGRSMYNNHNFHDFDRFHRFDYFMNFPDMFMDMSGYEMDFNGRWMNMHRF |
| 96 | Provided from MBL using Ref^4^ | Ocbimv22014011m_Scaffold212420_27387-27883 | *Octopus bimaculoides* | HPFHSFSGYHGYHHSGYYPYHSYSQGRRYHNYYDMFYDMSHYQMDFDGNWMDMYNHYSHPFFGYNHYHRGSHYYNHYPYHNYSWGHRFYDYPERFFDMSHYEMDFNGRWMNMHRF |
| 97 | Provided from MBL using Ref^4^ | Ocbimv22038094m_Scaffold88556_30839-37016 | *Octopus bimaculoides* | MLPSYVPDFDSQESQQLPPYLNRCNMLNGRNLSFPESMQQYDVSKRFAPQQFEMKRENVNRRTRNMPYFHDDDDDRTLLLTYDGGRILRPQSFHEIPTSCTEDKMNNQINFCSKRRSHAEPLESSLMTEDSKDLAENSLALIRVPRHDGEISLRQQTPLHSDMLHSQKNYPQQSNFSFVPRKSFLNPFLNPHLQRYNNFHKYTPNRPPFPTGFPNFINNPFRQNCPPFRMNPCGCNCYTQPTRFPTQEYCRFPTHRMKLSNPTRDYEWDFINDADDDNYHWNQNDVEPRTHFEFSNYQRRKHQNNFGLSQPIYCHDAISLQEISADPYSSNFNNETDIDKIEEEYFTYLQKTIPNFNVLPKKSNLKIASLIPEKKEKSLTNSTFASSNQTRNNFGDSHQTARSLMLMRLLKVLEENIDMSNYSMDFQENYVDSHGRLVDPFQTYITDLEENKLKSQIRDEAFVQETLSSRPPVVVLRNRRSDVITLDEGVTLKSSAKYMEKKEDDYGKNIKLSNASLHDYVNRSLDVPVRHQQTHQTQQQQQREEHPILQSHPEVIVETIRRERNVREDFSEPPSFTVPGVHERFFKTSVEQPIDMVTPWEKEREFQRYNMFKP |
| 98 | Provided from MBL using Ref^5^ | Adu_34322.1 | *Architeuthis dux* | NRYMMRHRPMFNSMSRTGRKYRGMMEPMSRMTMDFQGRYMDSQGRMVDPRFYEFYGRSNYDRSYGRPMFNSGPYMEGQRYGGWMDFPERHMDMSNYQMDMQGRWMDMQGRQCNPFNQWYNRPGYYPNYSYGRNMYYPERHMDMSNYQMDMQGRWMDNYGRYVNPFSHSMYGRNMQYPYNYNSSRYMDFPERYMDMSNYQMDMQGRWMDMQGRQCNPFNQSGYSRPSYYPGYSYGRNYYPERYMDMSNYPMFSSMSRSGRKYRGMMEPMSRMTMDFSGRYMDSQGRMVDPRFYDYYRNNEYERYYPRSMFNYRPYMEGQRYGGWMDFPERYMDMSNYQMDMQGRWMDMQGRYMDPWSNNYDTSNYMF |
| 99 | Provided from MBL using Ref^5^ | Adu_34324.1 | *Architeuthis dux* | NRYMNRFRHMFSNMSGNMYRGRNRGMMEPMSRMTMDFSGRYMDSMGRMVDPRYYDYYGRYDYDRYYGRSMFNYGWMMDGDRYNKYYRYMDFPERYMDMSGYQMDMYGRWMDMQGRHCNFNQWGYNRHGYFPGYSYGRHMFYPERWMDMSNYSMDMQGRYMDRWGRHCNPFSQYMNYYRYWNYPGYSNYYYNRYMYNPERYFDMSNWQMDTQGRWMDMQGRHMNPYWYNSYGRQMYYYQNYWYGRFDYPGMDYSNWQMDMQGRWMDMQGRYMDPMMGDSYYNNCNVLTTQAVHIYILHCQSGSVPYSSGETGKYCGCFLYPPIYFLKDCEGGPGVGEHERHQVHSRRL |
| 100 | Provided from MBL using Ref^5^ | Adu_34315.1 | *Architeuthis dux* | LHDLTFPRCSDFIMNRYMNRVRNWYGNNYRGRYRGMMEPMSRMTMDFSGRYMDSQGRMVPRYYDYYGRYYDYDRYYGRSMFNYGWMMDGDRYNRYYRYSDFPERYMDMSGYQMDMYGRMDMQGRHCNPFNQWGYNRHGYFPGYSYGRHMFYPERWMDMSNYSMDMQGRYMDRWGRHCPFSQYYNHWSRYGNNSSYYNYYYMYYPERYYDMSNWQMDMQGRWMDMQGRHMNPYWFNWGRQMYYPYQNYYWYGRFDYPGMDYSNWQMDMQGRWMDMQGRYMDPYWMNDYYNNYYYYY |
| 101 | Provided from MBL using Ref^5^ | Adu_34316.1 | *Architeuthis dux* | FHAIKNVLTATYEIIMNRYMNRYRPMFNNMFSNMFRGRSRGMMEPMSRMTMDFSGRYMDQGRMVDPRYYDYYGRYYDYDRYYGRSMFNYGWMMDGDRYNKYYRYMDFPERYMDMSGYQDMYGRWMDMQGRHCNPFNQWGYNRHGYFPGYSYGRHMFYPERWMDMSNYSMDMQGRYMDWGRHCNPFSQHMNYYGRYWNYPGYSNYYYYNRYMYNPERYFDMSNWQMDTQGRWMDMQGHMNPYWYNSYGRQMYYPYQSNQWYGRGGDYPGMDFSNWQMDMQGRWMDMQGRYMEPHWNDYYSKYYY |
| 102 | Provided from MBL using Ref^5^ | Adu_34314.1 | *Architeuthis dux* | NRYMNRYRPMFNNMFSNMFRGRSRGMMEPMSRMTMDFSGRYMDSQGRMVDPRYYDYYGRYDYDRYYGRSMFNYGWMMDGDRYNKYYRYMDFPERYMDMSGYQMDMYGRWMDMQGRHCNFNQWGYNRHGYFPGYSYGRHMFYPERWMDMSNYSMDMQGRYMDRWGRHCNPFSQHMNYYRYWNYPGYSNYYYYNRYMYNPERYFDMSNWQMDTQGRWMDMQGRHMNPYWYNSYGRQMYPYQSNQWYGRGGDYPGMDFSNWQMDMQGRWMDMQGRYMEPHWNNDYYSKYYY |
| 103 | Provided from MBL using Ref^5^ | Adu_34321.1 | *Architeuthis dux* | VNMYPGKYRGMMEPMSRMTMDFQGRYMDSQGRMVDPRFYDLYGKLKDYGRHYGRLMLNSPHMEGQRFGGWMNFPERHMDMSSYQMDMQGRWMDMQGRYCNPFNPWGQNRSNKYQGALEMSRMSMDFQGRFMDSQGRDTILGNKTFLVTGAMGADIMVVFLTTGCLILVYI |
| 104 | Provided from MBL using Ref^5^ | pred2_20770.1 | *Architeuthis dux* | TNHDKSGGFTKNTKVDSNTPGTSAVTRASGDQSKTKGKQPSLLHIDRDLRFRRHMIPKRPMFAMDGPNRMMAPHSRDINRTRGPNFMLRDRRPSMMSARNMFKDAGAQMVPPPPRGFMPMQGHPKEPKDMQRDFTKPPIDGERRMPDFPQRQDNMSKYTMDFPGRMMDYPGKRVEPFKGPHPMPAQPKQESFARYPDRQLGMPTFGMNKPYNQQGNPMGRVPHAPSRYMMSMPPSYKIIRFPERYMDLSDYTMDFQGRFMNQQGKYVNPLNRARIHQGRVMMPVPPPIKQRFMQKERYMDMSRYTMDFDGRYRDNYGRQIDPMEQFNANVGRYIRNRPPSNRRLARFPERYMDLRYTMDFQGRFMDTHGRHVDPLGRFHRNYGKMPVNPPQNQRRFTRHPEKYMDMSGYTMDFGRFLDRHGRPIDLLGRHLSHPGRFMLPSPDMQRDVSAIPRRFVAPPLKQIDMSRCSMDFGRLMDNQGRYVDQMRRFGDHMIPAIPRRFVAPPLKQIDMSRYSMDFEGRLMDNQGRYVDMRRFGEHLVPEPQINQGNFMDFSRYTMDFQDHLAGKYNSLVDPVGRLPALGPKPNIFFERSSPELRPTSVPENVPPMEPDPIMPEKDYTELKPRPEFHRFRSPGEHRVEFQEPPHQESSPTPPDEETVMLLEDEHMQPPFMKLPMSPSMYSGMRSSMFSMPEDCFQMVPPMMHFNYEFMPGQQPSADNMMMMEMMNTGYPLYQSRRASFVAPESDYIMNEQQWFMGDCPECMDNEMHRNDRMTMMMPPGTHNRSSFRMANGPRQQGHLPSVMEGAADETRHEDQRSSRFDDRYSHMEDKEFDFRHVRFGYNEDYPGLMPEVHQTATNGLLSYGDHDDYMFDGGYYPGYHMDDFYEGFMPSPPMGFMNDFQSPFCFDPEFMGMHPGFIDEAGSYPYPCMSDNFGFGYYDDDEGELDQGEEEEEEGEEEEERRHRMRKHCERNRTPSPMRSQFHPHSPRRRSPSPHQDGEDIEATHRGMTRPRSFLREEEHFFDDKHARYLEEAAECLRERGRFLREQHERFLQEMEDTGPLSPQIQAEAEAHAHMVEEEVNSTEAEAEAVERLARTARRVSNAENQCALFGNRHHLVDSQSYSHKHMDEH* |
| 105 | Provided from MBL using Ref^5^ | pred2_34310.1 | *Architeuthis dux* | HQLFWGLMEPMSRMTMDFEGRYLDSQGRLVDPRYYRNYKYKANYPNSYDWYGSAANRYPQTDRCVYETNQYDRPWQTDDYESRLDLYTGMNWDFHDNFER* |
| 106 | Provided from MBL using Ref^5^ | pred2_34325.1 | *Architeuthis dux* | NGDASIVKKAELSTQESPECQPVYDGNMESMSKMTMDMKGRYIDSQGRLVDPTRAPDGKPNDQGIRSVGKRSVDPESRPSDMPTNTMDFGGHNGQGMPFDPYMDRGYMGHMEPMSMPYAPERFGVHMRNPERYMDMSRYTMDFEGRYMDPNGQFFDPYGRRSRTSRMMGGDNPIPSQTEITFGFVDKWYPEEWGAKLVSRQGNFITVQIHSGCDCLVGVWDFILETMTFDKGTYCFQFDPIYILFNPWCKGDQVYMDDKDLLQEYILNDHGHIFQGSGSSSVYQKPWNYGQVS* |
| 107 | Provided from MBL using Ref^5^ | pred2_34311.1 | *Architeuthis dux* | NRFQDSKSFMGFHSPSHFNRNDKYPGVMEPMSRMSMDFQGRFLDSQGRLLDPKVSEFYRYNFGGQNVPGHGHGGQHHGGYPYYMFPYFGGYMRGDSYPRFGRQMNFPERSMDLSNYQMMQGRWMDMQGRYTNPFNYQAGFYGRPNFHGWHPYMPNHWRFMHRPERHMDMSNYQMDMQRYMDMQGRYCNPFNQWGHHKSDKYPGAMEPMSRMSMDFQGRFMDSQGRLLDPKVSEFYRYNSGGQNVPGYGHGGQHHGGYPYFMSPYFGGYMRGDSYPRFGRQMNFPERSMDLSNYQMMQGRWMDMQGRHVNPFSGYYSRQRNHPGFSYVRIMYYPERYMDMSNYQMDMQGRWMDMQRNVNPFSGYYSRYMNHPGGYYFFSRYMNYPERSMDLSNYQMDMQGRWMDMQGRHVNPFNYNRQGYYPGSFYGRHMFNPERYMDMSGGQRNTFGDEFGSSPGRHAGEPQQVLGMANMDMGPHMRNMDMHSPHMRNMDMQGPHMRHMDMQGPQMRNMDMQGPHMRNMDMQGPHMRHMDMGPQMRNMDMQGPNMKNVDMHGPHMRNMDMQASHMRHMDMQGPHMRHMDMQGPQMRNMDMVPHMKNMDMQGPHMRNMDMQAPHMRNMDDINQYHHQHGCHHQPNMFNHHGYGNYGSNMMSGYEY* |
| 108 | Provided from MBL using Ref^5^ | pred2_34323.1 | *Architeuthis dux* | SQFETRFDIRLCQRRITS*SAIMNRYMMRHRPMFSTMSRTGRKYRGMMEPMSRMTMDFQRYMDSQGRTVDPRYYESYGRDNDYHHYYPRSMFNYRPYIEAQRYSGWMDFPERYMDMSNQMDMQGRWMDMHGRHMDTV*ISAQLTEMCVIETLLKNSSLKRDFEGRKSIINHGRPMPKPNLRSSNKKCDWLTGNAQLNKLFCWPCLLFNLEKGQFGMIKVTMIFNNLHMAMLKHERSRHIKFIIDLKTFGNLLQRSNNMISRYVQQSNNMLSCRVQQSNNRLSCRVQQSNNILSCRQQSNNMLSCCVQQSNNMLNRRVQQSNNMLSNKMATFKSVSQRKRKIRRYVCYNRSTI* |
| 109 | Provided from MBL using Ref^5^ | pred2_34312.1 | *Architeuthis dux* | HQLFWGLMEPMSRMTMDFEGRYLDSQGRLVDPRYYRNYKYKANYPNSYDWYGSAANRYPQTDRCVYETNQYDRPWQTDDYESRLDLYTGMNWDFHDNFER* |
| 110 | Provided from MBL using Ref^5^ | pred2_34313.1 | *Architeuthis dux* | KGVMNKLSTTFGKKPRETKKTPAEVKTRPSTFPVTSRIYTGTSIRPNFPLSSRKDSQIPPGTSSTLAKETSTQVSKEGAGYSRRLHENKTTKTSRTDDSKLYRDSEEDKIHDNKTIKTQTKDSKLYRDAVEDGIHKDKTGTQLQTKDSKLYRDSVEDGIHKDKTEKKLQTKDSKLYGFVEDILIKETRPPPNTPQKLRHPRIDSDNRTDVGKKHRDEEQADDDDFTNKNVSGLSDDGEYRTSNKNQSGDPQAHNDSDKDEKYGVTLVKALQFFQDESDDDIDGGKSSDCGPVERHDFPIEKDKNETEDSQASASPFIEFPDSDKDISLDIKRNIVYLLPKSDPPSPEDPHVTGSFKTDTDRSIKSDDYLSERYDRISHNPEQVLSGYSEGVYLRDISKEDKNNFSTQQEKSTLCDQDPVNYIHKNNNKKSSSDILEDVDPCRPINNHTTSKYNYPSASRSGNHLEGGEMFRRYEDLFMDTDGLDGYREGAYCIYGSNEDIGNICAQHEPSTSTYDQHPIRNEYDDDTFHPDKKHVNPFRLTNDQALRKYYYESASGHHFEREKMVRQNPEEFFVEHEGNPDRVSGDNSAEALVINQRSKEETGRINPKQHLPNHSQDLASSFNKDIDSNKDDGRMFKDDEKRLPMKSDTVPEGHKAESDNLQTSMDTAANLSQSSEGSSNRQLSLFGRWMVTHGQLVDPHNFCLWSYAYPDLYIDLSKYQMDLQGRWLDTKGRYITPFFYCYSYDGEAVYLGDCDYYDVFNNSYDIDDYSNEFVNNPERYIDLSSFQMDMEERWMDCGEYIDPITDWYDIDSEEDEYYEYGSLCINEAHDLSGLLDATDGYYQDFSHEDNLYLPKANELVSDRYYTDYPSAGDYIFWDYPAGYGDTDVAASMQNLYLGSGYFCDDFKGTNKEDFDDKDEPCSYHSGQYLPKNKTK* |
| 111 | Provided from MBL using Ref^5^ | pred2_34307.1 | *Architeuthis dux* | DPMNYEGLSQPRHKSGDLSQNCMRSFHKSQRDMMRRDITAKSNKNRRFGDLLEPMSRLTDFHGRLIDSQGRLVDPSRYFDMDDHYMDNDRFIYPHDVIRKPQGMHGYMQGGNAYGSLGNSTHRGKFTDGMYRDMHNGMHGGMQCSGYMQGGSMQNRPMMDGLQMYMQGGYMGDPYYYNYNPGMQDPSVDMHTYYNDQEGRQGXMRTLKSNLEPKRYHMKQTVV* |
| 112 | Provided from MBL using Ref^6^ | OctVul6B008574P1 | *Octopus vulgaris* | MYGHGYNGRMEPMHMMSMDFQGRYMDSYGRMVDPRSYGFYGRYHDQDRYYGRSMYNYHGFHDFNRFHRFDYFMDFPDMFMDMSGYQMDMYGRWMDMHGHYSSPYSYMFNSSRHGYNTSYYNGRYGFYPERFMDMSHYQMDMYGRYMDRNGRHCSPYNNYYRRYMNYPYMNFLHMYHPERSMDMSHYQMDFNGHWMDMYNHYSHPFFGHNHYYRGSHYHNYYPHSNYSWGHRFYNYPERYFDMSHYQMDFNGHWMNMYGHGYQPFHSFNGSNAYHHSGNSYSWGHRYHNYYDMFNDMSHYQMDFDGNWMDMYNQYSYPFFGYNHYQRGSHYYNYYPHYNYNWGHRFHDYPERFFDMSHYEMDFNGRWMNMHRF |
| 113 | Provided from MBL using Ref^6^ | OctVul6B013700P1 | *Octopus vulgaris* | MNRYMSRRNNFSRRYRGIMEPMSRMTMDFQGRYMDSYGRMVDPRCYSFYGRYSDNDRYYGRSMYNYYGFYDNDRFHRYGNFMDFPERFMDMSGYQMDMYGRWMDMQGHYSSPHWQMFNSSRHGYYPGYHYGRNWFYPERFMDMSHYQMDMNGRYMDRYGRHCSPYYNYYRRYMNCPFMNFLHMYYPERFMDMSGYQMDMNGRWMDNYGRHSSPFYSYHGRYYHNYPYYNYSWGQRYYNYPERYFDMGNYQMDFDGRWMDTYSRHCTPFYNYHGRFQHNSPYHSYSWGQRYYNNPERYFDMGNYQMDFDGRWMDNYGRYSSPFNSYHGKFQHNYPYYNYSWGQRYYNYPERYFDMGNYQMDFEGRWMDNYGRYSSPFYNYHGRYHHNYPYYNYSWGQRYYNYPERYFDMGNYQMDFEGRWMDNYGRNFHGYNYHNRHYYNSYPNYYNYNWGQRYYDYPERNFDMFNNQMDFDGRWMDGQNFYSSWDNYNY |
| 114 | Provided from MBL using Ref^6^ | OctVul6B013700P2 | *Octopus vulgaris* | MRKQKLRLFFSLSLLISSMNRYMSRRNNFSRRYRGIMEPMSRMTMDFQGRYMDSYGRMVDPRCYSFYGRYSDNDRYYGRSMYNYYGFYDNDRFHRYGNFMDFPERFMDMSGYQMDMYGRWMDMQGHYSSPHWQMFNSSRHGYYPGYHYGRNWFYPERFMDMSHYQMDMNGRYMDRYGRHCSPYYNYYRRYMNCPFMNFLHMYYPERFMDMSGYQMDMNGRWMDNYGRHSSPFYSYHGRYYHNYPYYNYSWGQRYYNYPERYFDMGNYQMDFDGRWMDTYSRHCTPFYNYHGRFQHNSPYHSYSWGQRYYNNPERYFDMGNYQMDFDGRWMDNYGRYSSPFNSYHGKFQHNYPYYNYSWGQRYYNYPERYFDMGNYQMDFEGRWMDNYGRYSSPFYNYHGRYHHNYPYYNYSWGQRYYNYPERYFDMGNYQMDFEGRWMDNYGRNFHGYNYHNRHYYNSYPNYYNYNWGQRYYDYPERNFDMFNNQMDFDGRWMDGQNFYSSWDNYNY |
| 115 | Provided from MBL using Ref^6^ | OctVul6B014527P1 | *Octopus vulgaris* | MNRFMNRFRPQFNRKYRGFMEPMNMMSMDFQGRYMDSYGRMVDPKQYEFYGKYSDNDRYYGKSMYNYYGFYDNDRFHRYGNFMDFPERFMDMSGYQMDMNGKWMDTQGQNSNPYWNMFSSSRQGYNPGYSYGRNWFFPERFMDMSHYQMDMGGRYMDKSGRHCNPYYSHYRRYMSHPQMNFSQMHYPERFMDMSSYQMDIGGRYMDKWGCHINPFSTYYFGKQSYYPHNFWSQRKYMDMSFNQMDMQFNSMDMNSRNCDQLHYFRNFDMWNNQMDFDGHWMNMNNQSYQPPSFIRSQVYYNPYHFYTWMSRYYSQPEKFYDTTNYQVEFGGKWPSQEYECITQE |
| 116 | Provided from MBL using Ref^6^ | OctVul6B021549P1 | *Octopus vulgaris* | MLNGRKLSFPESMEQHDVSKRFAPQQFEMKRENVNRRTRNMPYFHDDDDVDRTLLLTYDGGRLMRPQSFHEIPGSCTENKMNNQINFCSKRRSHAEPLESSLMTEDNKDLADNSLALIRVPRHDGEISLRQQTPLHSDILHSQKNYPQQSNFPYVPRKPFLNPYLNPHLQRYNNFHKYTPNRPPFSTGFPNFINNPFRQNCPPFRMNPSGCNCYAQPTRFPTQEYCRFPGHRMKLSNPTRDYEWDFINDADDDHYQWNQNDMDPRTHFEFSNYQRRKHQNNFGLGQPIYCHDAISLQEISADPYRSHFNNETDIDEIEEEYFTYLQKTIPNFNVLPKKSNLKITSLIPEKEEKSLTNSTFASSNRTRNNFGDPQQTARSLMLMRLLKVLEENIDMSNYSMDFQENYVDSHGRLVDPFQTYITDLEQNKLKSQRDEAFVQETFSSRPPVVVLRNRRSDVITLDEGVTLKSSAKYMEKPENDYGKSIKLNNASLHDYLNRSLDVPVRHQQAHQTQQQQREEQEKHPILQSHPEVIVETIRRERSVREDFSQPPSFPVPGVHERLFKTSVEQPIDMGTPWEKEREFQRYNMFKPQFVPETRIYSNAAQIPLSHDIFDGETGKRDVINISQQRLIRKNGDPNEIPYETMAEYFQSSAADINPRVSTKYVFLKDPNSEGIQEDILNKLKCQSKLRSFDEKPFVTEAPDESDPENEVPDRESEFLGRQSFMALKQRFETEKHVQFVDEDNNGSFKNYPYFKSRVDSSDYSY |
| 117 | Provided from MBL using Ref^6^ | OctVul6B022188P1 | *Octopus vulgaris* | MNRSRNMFRNSSRKHRGVMEPMTRMTMDFQGKYLDSSGRLVEPRCNDYYGRNSNYRSMHNTGFYDNDKFQKHGRFMQFPERHMDMSGYQMDMRGRYMDKYGRHCNPYSRRHMNYPSNNYENYHMYNPEKLMDMSNFQMDMHGRWMDSNGRYSSPFSNYGSRHHQNYPHVNYSWGQRGVNYPDRFFDMSNYQMDLDGKWMDTYGRHCHPFYDNSNYYGKQYSYNMYPHYNYNWGQKYYHYPERYFDMSNYQMDFDGRWMDMFGRNQSPFNGYNNNNQGRQHHGQPQNSFSYGQRYQDRNFDIGNYQMDFDGRWMDMYGRYSNPFYGYNNFHGRYQHSLPHHFNWGQRYTQHPERLFDMGNYQMDFDGHWMDMDDRQCQPFSGNYNHSNRYQQNSNPSQSHNFNWGQRYQDYPERFFDMSGYQMDFDGRWMDSNNYSSDNFW |
| 118 | Provided from MBL using Ref^6^ | >OctVul6B023351P1 | *Octopus vulgaris* | MDKRHQQGSTDRLKHWPFLNYFNRSRNMPPVCPPPDTSSKDKKDPMGRMPRGNVRHFGPPFMGPPMPGEGFEGFPPPPPPCAMSAPPQSRKKKNAEKERLQDVEGEPRYEDNTEKRPKSPAATQERKSEAKDKDVREPRQENHHNRNDPCDKQSDVGSQHRNSSVFDTESKFSDLEKSNRFYPYPEFDRHGMMPPFDMMPMGYGPPDGMRPPVYMNYPEKPVDMSRMTMDFEGNYMDNNGQFMDPLDMPQHNEMDLLRENMNFRPPYMDQFPGDMDGCVGIPPMEMKRYTQRRKDGFDEMSPQELMENGMLIVTHVDYHLDETLRLHHTDQYDIGQNGQLIVRRGQPFMLTIEFHEEYNEEKHKIRFIFQIGEHPLPSKKSDIRFGLTDKWYPEEWGAKLVSRRGRMITVCVHPACDCIVGVWDFMIKTTVSGKGSYIFDNFEQIYILFNPWGQTDQVYMEDKDLLYEYVLNDHGHIFQGSGCSSVYQKPWNYGQFDEDILDIALYLMRQGFLEYCPQMSSPVRVARVLVHVINSPENTGVMMRFDSDNCQNGKKPTAWGGSRRILQQFIDTKEPVKYGQCWVFSGLLTTVCRALGIPCRTVTNFNSNHDSDDNLSVNVYLGEDEKGNIFEQKEGNVWNFHVWNDVWMSRSDLPVGYCGWQAVDPTPQEASDGIYCCGPAPLKAIKNGEVNQTFDTNFIFSELNADRVYWKKNHRSGKWEIVHIDRNALGKFIYTKYPNCKPGHNRSGGLMDITFDYKLVNGSEYDRIDVINVHRKKMMLHRAKKSSHYQEDVEFRISERESVMVGQDFNITLHCRNTGQDQRYVSSSVYCKIVDHFGEKIGICRELHVNNFFDAKNSKMIVMNVAPEDYMPFLGRDNNSRLSMRIMVYCKVKQTDQMFVFNDSYQLDWPYLTIEVPPKIRVRDEVMARISFVNPLDIPLTNCELMLEGNGFERIYEIRVSDVPPHGNFMEEIVFIGRKIGEKQLVATFYCQELCDIVGSSMLQVFK |
| 119 | Provided from MBL using Ref^6^ | OctVul6B023351P2 | *Octopus vulgaris* | MYTILKYFSKKMSNKTMPAEDTSSKDKKDPMGRMPRGNVRHFGPPFMGPPMPGEGFEGFPPPPPPCAMSAPPQSRKKKNAEKERLQDVEGEPRYEDNTEKRPKSPAATQERKSEAKDKDVREPRQENHHNRNDPCDKQSDVGSQHRNSSVFDTESKFSDLEKSNRFYPYPEFDRHGMMPPFDMMPMGYGPPDGMRPPVYMNYPEKPVDMSRMTMDFEGNYMDNNGQFMDPLDMPQHNEMDLLRENMNFRPPYMDQFPGDMDGCVGIPPMEMKRYTQRRKDGFDEMSPQELMENGMLIVTHVDYHLDETLRLHHTDQYDIGQNGQLIVRRGQPFMLTIEFHEEYNEEKHKIRFIFQIGEHPLPSKKSDIRFGLTDKWYPEEWGAKLVSRRGRMITVCVHPACDCIVGVWDFMIKTTVSGKGSYIFDNFEQIYILFNPWGQTDQVYMEDKDLLYEYVLNDHGHIFQGSGCSSVYQKPWNYGQFDEDILDIALYLMRQGFLEYCPQMSSPVRVARVLVHVINSPENTGVMMRFDSDNCQNGKKPTAWGGSRRILQQFIDTKEPVKYGQCWVFSGLLTTVCRALGIPCRTVTNFNSNHDSDDNLSVNVYLGEDEKGNIFEQKEGNVWNFHVWNDVWMSRSDLPVGYCGWQAVDPTPQEASDGIYCCGPAPLKAIKNGEVNQTFDTNFIFSELNADRVYWKKNHRSGKWEIVHIDRNALGKFIYTKYPNCKPGHNRSGGLMDITFDYKLVNGSEYDRIDVINVHRKKMMLHRAKKSSHYQEDVEFRISERESVMVGQDFNITLHCRNTGQDQRYVSSSVYCKIVDHFGEKIGICRELHVNNFFDAKNSKMIVMNVAPEDYMPFLGRDNNSRLSMRIMVYCKVKQTDQMFVFNDSYQLDWPYLTIEVPPKIRVRDEVMARISFVNPLDIPLTNCELMLEGNGFERIYEIRVSDVPPHGNFMEEIVFIGRKIGEKQLVATFYCQELCDIVGSSMLQVFK |
| 120 | Provided from MBL using Ref^6^ | >OctVul6B030895P1 | *Octopus vulgaris* | MYGHRYNGRMEPMHMMSMDFHGRYMDSYGRMVDPRSYGFDGRYHDQDRYYGRSMYNYHGFHDFDRFHRFDYFMDFPDMFMDMSGYQMDMYGRWMDMHGHYSSPYSYMFNSSRHGYNSSYYNGRYGFYPERYMDMSHYQMDMNGRYMDRYGRHCSPYDNYYRRYMNYPYMNSFHMYHPERFMDMSHYQMDMNGQYMDRYGRHCSPYDNYYRRYMNYPYMNFLHMYHPERSMDMSHYQMDFNGHWMDMYNHYSHPFFGHNHYYRGSHYHNYYPHSNYSWGHRFYNYPERYFDMSHYQMDFNGHWMNMYGHGYQPFHSFNGSNGYHHSGNSYSWGHRYHNYYDMFNDMSHYQMDFDGNWMDMYNQYSYPFFGYNHYHRGSHYYNYYPHYNYNWGHRFHDYPERFFDMSHYEMDFNGRWMNMHRF |
| 121 | Provided from MBL using Ref^7^ | Ebe_EB43639 | *Euprymna berryi* | MFSVRYKDDYMVCIRFRHLPNVTTMNRYMNRFRNFYGNMYRGRYRGMMEPMSRMTMDFQGRYMDSQGRMVDPRFYNYYGRFNDYDRYYGRSMFNYGWMMDGDRYNRYNRYMDFPERYMDMSGYQMDMYGRWMDMQGRHCNPYSYWMMYNYNRHGYYPNYSYGRHMFYPERWMDMSNYSMDMYGRYMDRWGRYCNPFYQFYNHWNRYGNYPGYYSYYYMYYPERYFRHV |
| 122 | Provided from MBL using Ref^7^ | Ebe_EB43543 | *Euprymna berryi* | MNRYMNRFRNFYGNMYRNRNRGMMEPMSRMTMDFQGRYMDSQGRMVDPRYYDYYGRYNDYDRYYGKSMFNYGWMMDGDRYNRYNRWMDYPERYMDMSGYQMDMYGRWMDMQGRHCNPYSQWMMYNYNRHGYYPNYSYGRHMFYPERWMDMSNYSMDMYGRYMDRWGRYCNPFYHYYNHWNRHGNYPGYYSYYYMYYPERYFDMSNWQMDMQGRWMDMQGRYCSPYWYNWYGRQMYYPYQNYYWYGRYDYPGMDYSNWQMDMQGRWMDMQGRYMDPWWMNDSYYNNYYY |
| 123 | Provided from MBL using Ref^7^ | Ebe_EB43499 | *Euprymna berryi* | MTEDDGQRVQNQETEPVLKTTFFKITSTPVILFSFENHPLLEMNTFMDTMHCDGMGMPQSKFGDFSHNCMRSFPKSQRDLMRRDLVAKPGKNRRFGDFMEPMSRMTMDFYGRMIDSQGRIVDPNRFLFAEEHYMDNDRFPYFYDMMRSPRSMYSGMYGFGYGDHSFNRGIYNDDMYHGGMNPFMHNRSMMGRMYSPGPFMDDAFSMYYRPRMGDHFMYSQSQFNDHEGGQGMFGRMPDNFETSPGRPTEEQSIASRLSESHNLHRRLSENHARIEAANNQRKASRALIFPEETSNMESA |
| 124 | Provided from MBL using Ref^7^ | Ebe_EB43604 | *Euprymna berryi* | MALQTSRYYSTVDGVAALSATKELGPLYYQSETAARNCQSGTLNPSEEGQPNLIAQAYVALKHSQVPSFEGIGVRRLSRESNPSRFRLSSYPSLPRRFRCRSNVTTMNRYMNRFRNFYGNMYRGRYRGMMEPMSRMTMDFQGRYMDSQGRMVDPRYYDYYGRYNDYDRYYGRSMFNYGWMMDGDRYNRYNRWMDYPERYMDMSGYQMDMYGRWMDMQGRHCNPYSQWMMYNYNRHGYYPNYSYGRHMFYPERWMDMSNYSMDMYGRYMDRWGRYCNPFYHYYNHWNRHGNYPGYYSYYYMYYPERYFDMSNWQMDMQGRWMDMQGRYCSPYWYNWYGRQMYYPYQNYYWYGRYDYPGMDYSNYQMDMQGRYMDMQGRYMDYPYNYYNCY |
| 125 | Provided from MBL using Ref^7^ | Ebe_EB43274 | *Euprymna berryi* | MNRYMNRYRPMFNNMYGNMYRGRYRGMMEPMSRMTMDFQGRYMDSQGRMVDPRYYDYYGRFNDYDRYYGRSMFNYGWMMDGDRYNRYNRYMDFPERYMDMSGYQMDMYGRWMDMQGRYCNPYNQCGYNYNRYGYYPNYSYGRHMFYPERWMDMSGYQMDMQGRYMDRYGRYCNPFSQYMNYYGRFWNYPGYNSYYNRNMYYPERHFDMSNWQMDMQGRWMDNQGRYSSPYWNNFYGRQMYNPYQNYNSYGRCDYPGMDYSYCQMDGRCNDSGMGDSYYNNW |
| 126 | Provided from MBL using Ref^7^ | Ebe_EB43337 | *Euprymna berryi* | MNRIPDRKRFMPRQFYKSEKYRGELEPISMMTMDFQGRYLDSQGRMHDPRVYESYQRYHLSSPNYLNFDPSCSQSYTWGPYPGDYTKSDWYPRPRRQMVFPEKFMDLSSYQMDMKGRWMDTQGRYTNPFNSKNRARRNHFPLFSPQMYSNNYGNDMSHTEGNKDTPGYPIDSQDQWMYIQGRQTKPSSYDAVGFLGKHYPYKYPPYMHDTWRFMSWPERYMDMTGYQMDMHGRWMDTQGRHCNPFNQCGYNTQGSYLGDPYDRNIVYSEKLVDRSNDRIGRQERSAGRYGRRVNPLSRHSCRVCMDMNNPYPGKMDMSNYQMDMQGRWMDTKGRYTNPFPSSGYNKQGPFHSFQYKRNMPYPEKLIDMSNYQMDMQGRWMDTQGRYTNPFSVSGYNRQVPFSGFPYNRYMPYPEKMIDMSNYQMDMQGRWMDTQGRYANPFSFGGFTRQRPFHGFPYNRNMPYPEKMMDMSNYQMDMQGRWMDSQGRYTNPFSFSSFTRQGPFSGFPYNRNMPYPEKMMDMSNYQMDMQGRWMDTQGRYTNPFSFSGYNKQELFPGFSYYRNMPYPEKMMDMSNYQMDMQGRWMDTQGRYSDPFSFSSFNRQWPFLGFSYNHNTLHPENMMDMSNYQMDMDGHWMDTHGRYTKPLGFSSYRRQWPFSGFPHMLYPENMMDMSNYQMDMQGRWMDMHGRYTNPFSFSGFNRQRPFHSFPYNRNMSYPENMMDMSNYQMDMQGRWMDTQGRYRNPFNQYSNYNRYGYFPEYSFDRNTFPMRSMNISNNQIEVPGRFSEPYDHYLSTMDIVHPFYNYRYSRNIYYPDRYMDMSGNHVDMQEPLMDTESRFHPSYQQGRNMNPLDRLLDSYDYNSQTDMEGRWMDYYGEYTHPSFGDQGRKTFGDYFRSLPGTYHGDTYRYLGTRNIYDRDQFYYQHGYYRQPPVINNHNDGYYGNNMMEYIYDN |
| 127 | Provided from MBL using Ref^7^ | Ebe_EB43491 | *Euprymna berryi* | MSRQVKTGQGNNTLPQNSGNNNSQSLPHTSQTPPKGTHSMSQEGGESMPQGPQTMPREQPSMSHDPKSMKNKYPMMYEDYGGYSLPPPGSYPMGYGSPPMMSHRGFPMSPDMQSMQGEMYPMYGSPGSNSMSQANNKNRMKTLSKSQRENMFREGPKSGGSYQHSETSKPQTMDRMTMDYEGRFIDRRGRIVDYNRNMDSDSHRDDRFMFNRDMPRGDQGMYDYGMPAYMYGNMMESPDLMLDMDMDMDMYGRYPMFYNQYGDDDEEDYDDGMMDNMVNVSHGYMDGRYETDFAGLNYPDRRTMESVGRYGDSFER |
| 128 | Provided from MBL using Ref^7^ | Ebe_EB43237 | *Euprymna berryi* | MNRYFTRHRPMYSHMYGNKYRGMMEPMSRMTMDFQGRYMDSQGRIVDPRLYDFSGSRFNDHDRYYGKSMYGHGSYMDGQRYGGYMDNPERYMDMSNYQMDMYGRWMDMQGRYCSPFAQWSHNRQGNYPGNSYNRNMFHPDRRMDMSNYQMDMQGRWMDMQGRHCSPFNQMGHNRHGNYQWFWHSRYPERWMDMSGYQMDMEGRWMDNYGRYVNPFSNYSYNFGRGTNYPGSYNNYSFGRYMNYPERWMDMSGYQMDMQGPSMDMQGHYMDNFDRNYNDYQMF |
| 129 | Provided from MBL using Ref^7^ | Ebe_EB43298 | *Euprymna berryi* | MYRGRYRGMMEPMSRMTMDFQGRYMDSQGRMVDPRFYDYYGRFNDYDRYYGRSMFNYGWMMDGDRYNRYNRYMDFPERYMDMSGYQMDMSGRWMDMQGRYCNPYSYWMMYNYNRHGYYPNYSYGRHMFYPERWMDMSNYSMDMYGRYMDRWGRYCNPFYQFYNHWNRYGNYPGYYNYYYMYYPERYFDMSNWQMDMQGRWMDMQGRYCNPYWYNWYGRHMYYPYQNYYWYGRYDYPGMDYSNYQMDMQGRYMDMQGRYMDPWWMNDYYNYNYYY |
| 130 | Provided from MBL using Ref^7^ | Ebe_EB43586 | *Euprymna berryi* | MNRFMNRYRPMFNNMYSNMYRGRYRGMMEPMSRMTMDFQGRYMDSQGRMVDPRFYDYYGRFNDYDRYYGRSMFNYGWMMDGDRYNRYNRYMDYPERYMDMSGYQMDMSGRWMDMQGRYCNPYNQWGYNYNRHGYYPNYSYGRHMFYPERWMDMSGYQMDMQGRYMDRWGRYCNPFSQYMNYYGRYWNYPGYNNYYYSRNMYYPERYFDMSNWQMDMQGRWMDMQGRYCSPYWYNWYGRHMYYPYQNYYCYGRYDYPGMDYSNYPMDMQGRYMDQYGMNDYYY |
| 131 | Provided from MBL using Ref^7^ | Ebe_EB43580 | *Euprymna berryi* | MEKVISVRNGSSTFWREPRLAVRFKESAKDVLVENRAPVVSDSLLSQAFPEVVIIMNRYMNRFRNFYGNMYRGRYRGMMEPMSRMTMDFQGRYMDSQGRMVDPRFYDYYGRFNDYDRYYGRSMFNYGWMMDGDRYNRYNRYMDFPERYMDMSGYQMDMSGRWMDMQGRYCNPYNHGVTTTTDTVTIPTTPTAAICSTRRDGWTCLTTPWTCTDVTWTGGDVIATRSTNSTTTGTATATTPGYYSYYYMYYPERYFDMSNWQMDMQGRWMDMQGRYCSPYWYNWYGRHMYYPYQNYYWYGRYDYPGMDYSNYQMDMQGRWMDMQGRYMDYPYNYYNWY |
| 132 | Provided from MBL using Ref^7^ | Ebe_EB43342 | *Euprymna berryi* | MNRYMMKHRPMYNQMCRTGRRYRGVMEPMSRMTMDFQGRYMDSQGRMVDPRYYDFSGSSDRYSGKSMLNYGSYMDGGQRYGGFMDYPERYMDMSNYQMDMHGRWMDMQGRYNSPFSHYNYSYNRHGNYPGYYSYNRNMCNPERMMDMSNYQMDMQGRWMDNYGRHVNPFSHFMYGRNMHYPNFNYYSGRYMNYADMSNPQMDMQGRYMDSSMSNMYDNYNNYY |
| 133 | Provided from MBL using Ref^7^ | Ebe_EB10152 | *Euprymna berryi* | MIPKRHPMFPMNGPHRMMPPNARHANRMHSPNVMFRGRAPNMMPGRSMFQGPGAQMGMAPPGGFMAPMQHRTKESENMQPQNEGQGRMSGLTQRQEEMPKDSRGFPGKMMDFHTIPGQPKHGGPMRMHTGPMGMQSGLMGMRSGPMGMQSGPMGKQSGPMGMHSGPIGMHSGPMGMQSGPMGKQSGPMGMHSSPIGVQSGPMGKQSGPMGMHGGPMGMQGGPMGIRGSPMGKQSSPMRMYSGPMRMQSGPMRMRSGPMGMKGGPMGMQSGPMRMRSGPMGMHAFGMNKTSGQQGKPMGRFPRGLRRHMMPIPLSYKRMIRFPERYMDLSGYTMDFKGRFMNQQGKHVNPLHSARMHQGRAMMPVPLPIKRRFMQKPERYMDLSRYTMDFEGRYRDRYGRQIDPMEQFNANIGRYVRNLPPVHRRMVRFPERYMDLSRYTMDFQGRFMDSHGRHVDPFGRIHGNYGKVLMSPPLNQRRFTRQPEKYMDMSGYTMDFQGRFLDRRGRPVDLLGQHHRHPERFMPPVPYMQQPMPMMPRKFPVAPPFKPMDMSQYTMDFQGRLMDNQGQYVDHMRRFGDHMGPHPSHDQRNLMEFSRYPMDFQDRHFDKYSGPFDRVYKPNTSFERCGSPERGQSPGPETNIHIESRQIIPEKEYTELKSSPGFNRVNSEMSRPSSSGEQRVEMQYGPQQGNVLSAPSDMESYMFPPIDDEHMQMPFMPPPMFSPIHASMHQPMFSSEKPRMHYPTPEDDTQMQQHMVHFEDEMHMPMHKPFADDMMMMKMMEMSNAGYQQYQWMDDPFMFPESEYMMDDRQWHSDDGFDYMDDEMPYEYSQMPNMMAVGMHGRPTFRIDEETRDQSHLPPVIGSSDSSEIGRDERCDNRMDYKHMHSMEDREFDSRHGRFETHEDIDYPGFMQDMHQRPMSGFFDHCDSDDGMFDDDYYMDNNMDEDFFPNPRSSMAYPSMGFMNDFQMPFSFDPEYMGMQSGFMPEPEYYPYPYMDDYFGYGFDDDDSEDEREREEEEEEGRHRSMKHQTSSSLHRAFHSHGLYHGNMNTRQGVDDVATGIDRPSSLPRIVLSEDKPLVDDKHARYLEDAAEYLKERSRFLRDQHNRFVQELENAGPLKHPQVQAEAEAHANEVKEEVNATEAEAEAVERLARTARRLSNVDHQLAANRHPSFSA |
| 134 | Provided from MBL using Ref^7^ | Ebe_EB21986 | *Euprymna berryi* | MDDGGDRYNRYKPIMDFPEGTGHFWFQMECMGAGWTAGALLQTPTTNAGSITTDTVSIQIFSYGRHMFYPGVVDMSGFRWTSRALQEDMDAIANRSLDMNSTVDSGTTRVNS |
| 135 | Provided from MBL using Ref^7^ | Ebe_EB43404 | *Euprymna berryi* | MSKPQGQRTSHGLKSWPLFSYFNRSRGMPVASPPGPAKDHPAKSDAPMHMPVERQQNFNPPMMAQRAPRYSNPHIGPQGPGMHYDGSRGDASKQAHKGKKDATEKDRQRPIDNKQSRNDGRMESMSKMTMDPQGRYIDSKGKVVDPSTVSDGKRTGSQKRQNTPDDRVPNMPNNDMDFVGPYMEGQDRPYNPYMEPGYMGQMGPMPMRYMDPGHFRMPIDDPERYMDMSRFTMDFEGRYLDPSGQFFEPFGQQGDDPMMDMMTQQMHENNLGEEMTEEELIESGMLVVTDIDYHLPENVRHHHTDQYDLAQNGQLIVRRGQPFMMTIRFNEEYDETKHSLKFIFQIGDNPIPSKKSEVKFGFVDKWHPEEWGAKLVSRQDQFITVYIHTGCDCIVGVWDFLLETMTFGKGSYCFDQFDPIYILFNPWCKGDQVYMNDKDLLCEYILNDHGHIFQGSGSSSVYQKPWNYGQFDDDILDISCI |
| 136 | Provided from MBL using Ref^7^ | Ebe_EB43515 | *Euprymna berryi* | MSSLMDVRHYDGTCVPYQNFRYNYTRGFPKSHRDMVRRDLMVKSGKNRTFEDFMEVMSGMTMDFGGRMIDSQGRIVDPSFFDEYYIDYDRFPYFHDMMRSPRFMYSGMYGFGSGDHSFCRSMYNDDMYRDIYHGGMNPFMQNRSIMGRMYSPNRFIEDPFSMCYRPRMSYHFMDSQQEGGQGMFRSMSHNIERPTGRPTEAQSIARRLSASHNLQRRLSESHARIDAANNQRKTSRAMIFPEESSNMESD |
| 137 | Provided from MBL using Ref^7^ | Ebe_EB43451 | *Euprymna berryi* | MVYIYIDVFRWHRVSQSRTEHPSLQVVLSVCPGLPFSSRVLIDMNRYMNRFRNFYGNMYRGRYRGMMEPMSRMTMDFQGRYMDSQGRMVDPRYYDYYGRFNDYDRYYGRSMFNYGWMMDGDRYNRYNRYMDFPERYMDMSGYQMDMSGRWMDMQGRYCNPYSYWMMYNYNRHGYYPYYSYGRHMFYPERWMDMSNYSMDMYGRYMDRYGRYCNPFYQFYNYWNRYGNYPGYYNYYYMYYPERYFDMSNWQMDMQGRWMDMQGRHCNPYWYNWYGRQMYYPYQNYYWYGRYDYPGMDYSNWQMDMQGRWMDMQGRYMDFPFNYYNWY |
| 138 | Provided from MBL using Ref^7^ | Ebe_EB43590 | *Euprymna berryi* | MYRYMNRYQNMLIGHNGKYRSMAEQMSRMSIPPSERMMDPSYYDYYGNDHRYYRGSVYDRNGFWLGNEGHYWYDNWMDNPERYMDMSDYEMDMQGRWMDKHGRYCDPFNQWDCNMYYYNPYYSYGRNMFYPEIYMDMSKYQMDMEGRWMDKKGRYCDPFNDWGYNRHYYYPDYSYFNMLFPERWLDMSSYQMDMEGHWMDLYGRRVNPFSHWMDDGNIYCHQCGFYDDWCMDHPEDWMDMSGYQMDMQGRWMDSKGRYCNPFANFFDCYDMQYHGNNSFFGHLPGNRMSICRYPMDTHGQWMDNQERYDGDY |
| 139 | Provided from MBL using Ref^7^ | Ebe_EB43354 | *Euprymna berryi* | MYRGRYRGMMEPMSRMTMDFQGRYMDSQGRMVDPRYYDYYGRFNDYDRYYGRSMFNYGWMMDGDRYNRYNRYMDYPERYMDMSGYQMDMSGRWMDMQGRYCNPYNQWGYNYNRHGYYPNYSYGRHMFYPERWMDMSGYQMDMQGRYMDRWGRYCNPFSQYMNYYGRYWNYPGYNSYYNSRNMFYPERYFDMSNWQMDMQGRWMDNQGRYCSPYWNNWYGRQMNYPYQNNYFYGRYDYPGMDNYQMDMQGRYMDQYGMNDYCY |
| 140 | Provided from MBL using Ref^7^ | Ebe_EB43505 | *Euprymna berryi* | MKGVRSRLGLSLGKKPKETKTSLTDTKLQTSTMPGRSPLGTGLSSRTNLPLVAGNKRGQIPTRDTSRLVTNKAPLNPSKEGAGNPLKPNQKTASKNDYPKSGTNIIQDLLGTQTDSDSPNRIKEELGDNPQTGTGEFTGDYVSGLTEEMKDLNTNRENNDGVQMIRGTTDSDKAGDSLDKHGVSIVKSIQVIQDEDEDDENTDKSSSSSSGKRVDFPTNDDKNTTADPKRSGSPFIEFPDGDKDAPRPDREDIVYMYPKGTPPSPEVKRVAAESQSRVDMESTERSNAYFCAEDERLSESPELNLDDRSGEVYFNHKLKKDYSDLSKPYNENTSMYVEDPINYDEFKKNRKFDSDSPEAVRSFRSGYDPKTYQLSGYDRKPINQLSKNRFDSDFGDDSDSNRLGNKRILSDSGEFRKNTHGFGTDNEGPYSDYGPNEESRLPNMLDAPTTLNDQPPTRNKYENDKTSVNPRNLDGEESMRLYEYEFGGHCQSGRNPSQDFEGFTEERENSPERISKIDYIEKKVIQRASARPVQENLQHSQKSQKSDNSQNSSTDFAKNKEANKREARMSQNNNAKDLHSKGDYDSQTDQNTLKDRNIYSTLKNEDGSSRQLSIFGRWMVTHGNLVDPQNFCPLAYATQPDLYINLSNYEMDAHGRWLDKNGRYVTPFFYCFGYDGEPVYLGDCDYTNGVHDNYMNDNYNDDFVNNPERYIDLSDYQMNMDKQWVNSLGRDIDPLSDWYSDGSEDDEYNEYSWMYDRPQFLEDSSLPNPMDGRYPNFCQPDTPYFNDADDRFPMGEAFSETCEYWDFAGCMENGIESMQHPWQAAGDFSRGYTGANEEYFVEEPDSVPYQQRSSKDSDK |
| 141 | Provided from MBL using Ref^7^ | Ebe_EB43501 | *Euprymna berryi* | MCRTGRRYRGVMEPMSRMTMDFQGRYMDSQGRMVDPRYYDFSGSSDRYSGKSMLNYGSYMDGGQRYGGFMDYPERYMDMSNYQMDMHGRWMDMQGRYNSPFSHYNYSYNRHGNYPGYYSYNRNMCNPERMMDMSNYQMDMQGRWMDNYGRHVNPFSHFMYGRNMHYPNFNYYSGRYMDYADMSNPQMDMQGRYMDSSMSNMYDNFNNYY |

**Table S2.** Identity matrix of reflectin protein sequences within individual species. For each species, the highest and lowest percent identity values of reflectin protein sequences were calculated from a multiple sequence alignment using Clustal Omega.

| **Species** | **Highest Percent Identity (%)** | **Lowest Percent Identity (%)** |
| --- | --- | --- |
| *A. dux* | 99.7 | 0.7 |
| *D. opalescens* | 74.7 | 27.4 |
| *D. pealeii* | 98.8 | 15.4 |
| *E. berryi* | 99.0 | 8.1 |
| *E. scolopes* | 99.7 | 5.8 |
| *S. officinalis* | 99.7 | 23.1 |
| *S. pharaonis* | 97.5 | 57.7 |
| *A. argo* | 92.5 | 18.4 |
| *O. bimaculoides* | 98.8 | 21.3 |
| *O. vulgaris* | 97.0 | 16.1 |

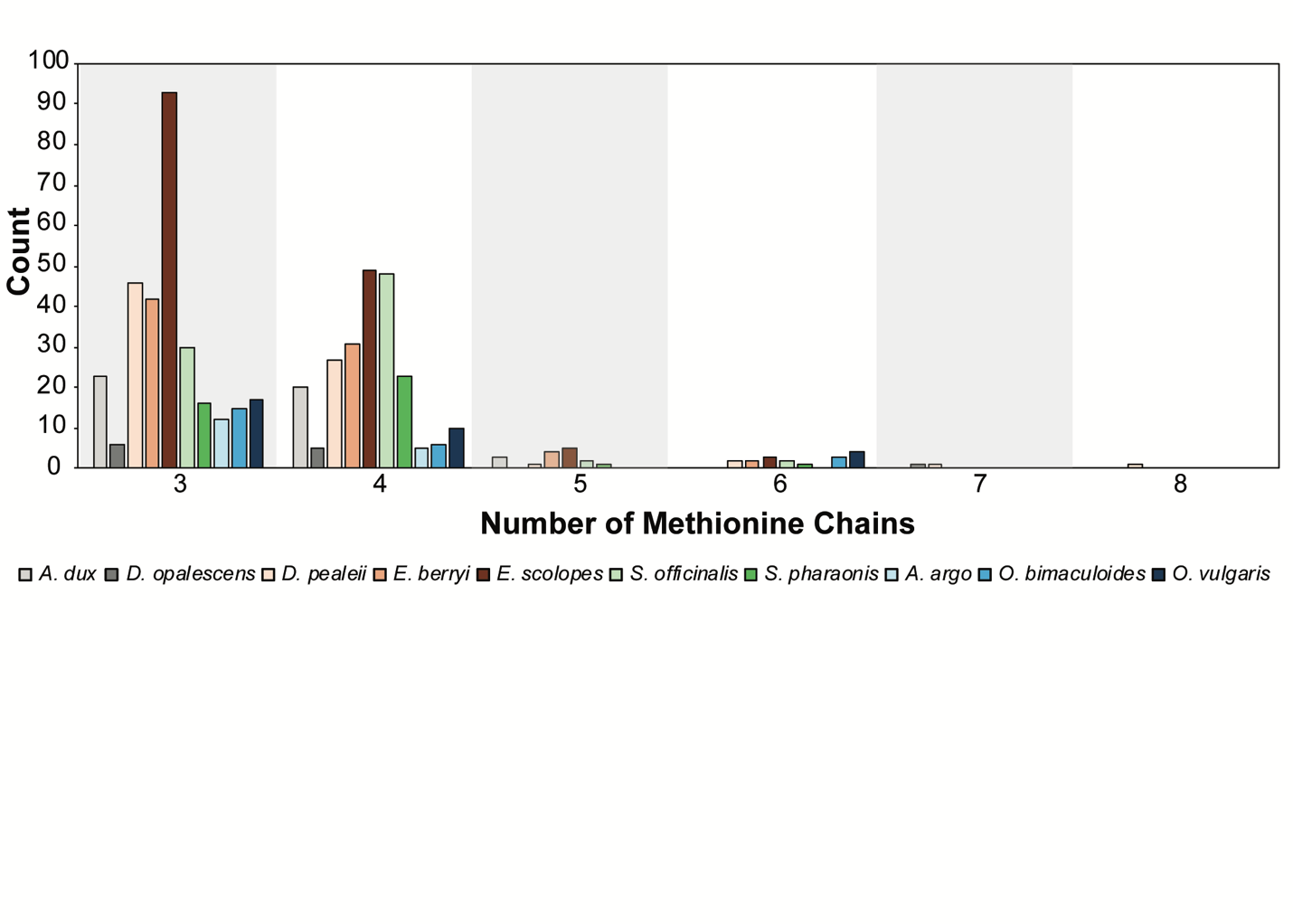

**Figure S1.** Stratification of reflection domains based on the number of [MZ(X)_5_] repeats. 141 sequences from ten different organisms were analyzed for the characteristic methionine-repeating sequence previously described. After identification of 560 reflectin domains, each were separated based on number of methionine chain repeats. Each different colored bar represents a different species.

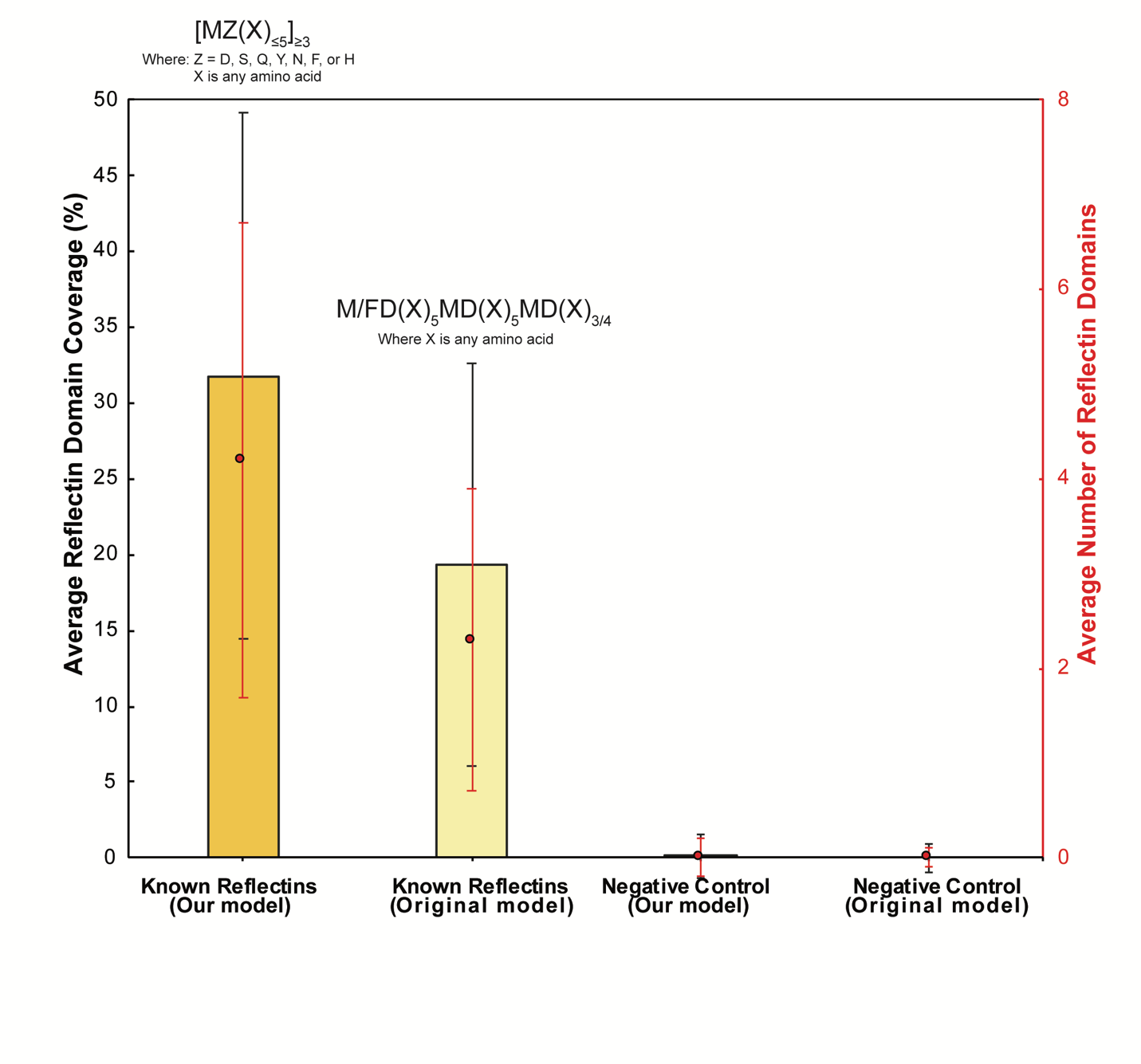

**Figure S2.** Comparison of the domain coverage between the original and our relaxed definition. The bar represents average reflectin domain coverage as a percentage with the red dots representing average number of reflectin domains compared between original to relaxed model for both experimental and negative control.

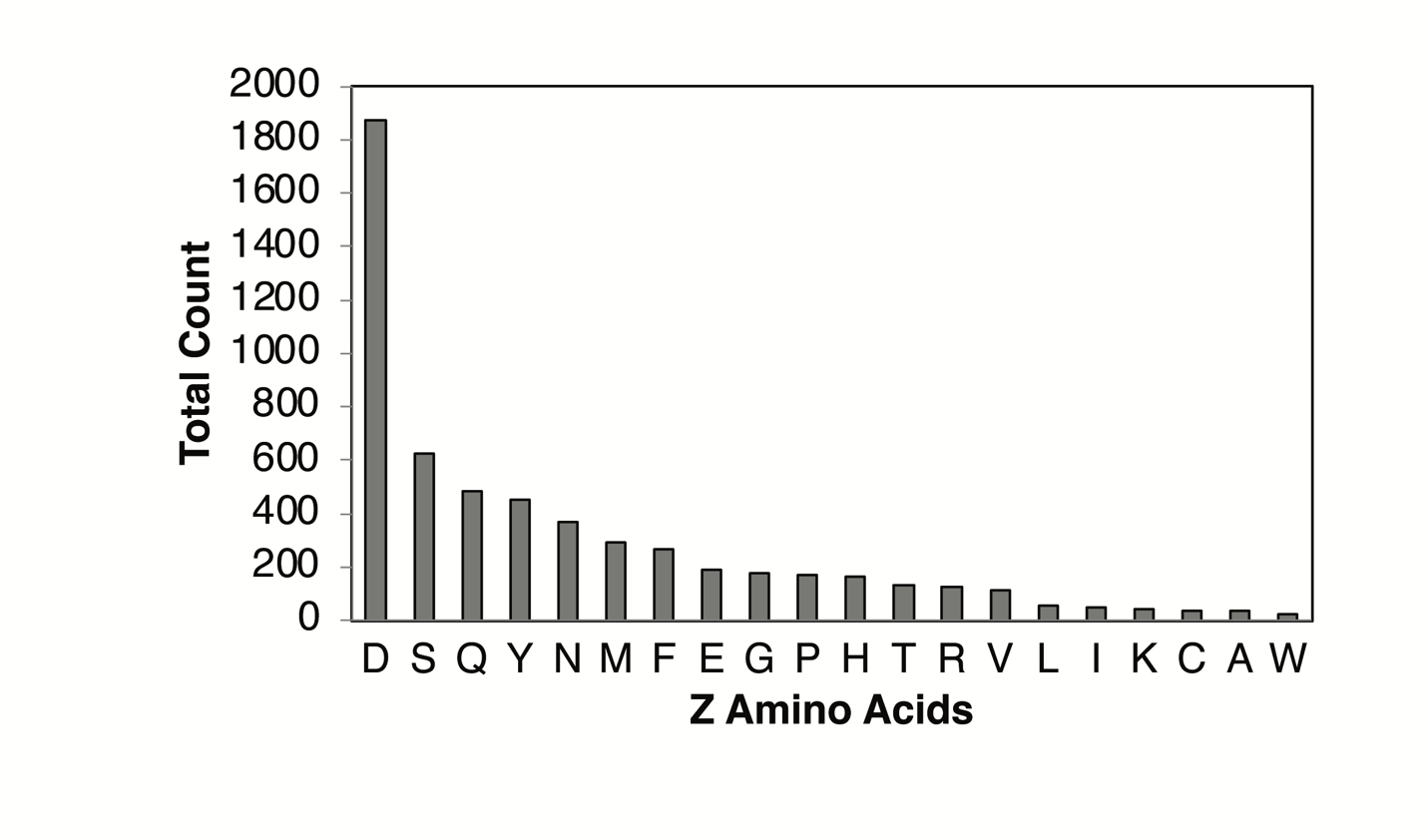

**Figure S3.** Total count of residues apparent in the “Z” position across all 141 reflectin sequences.

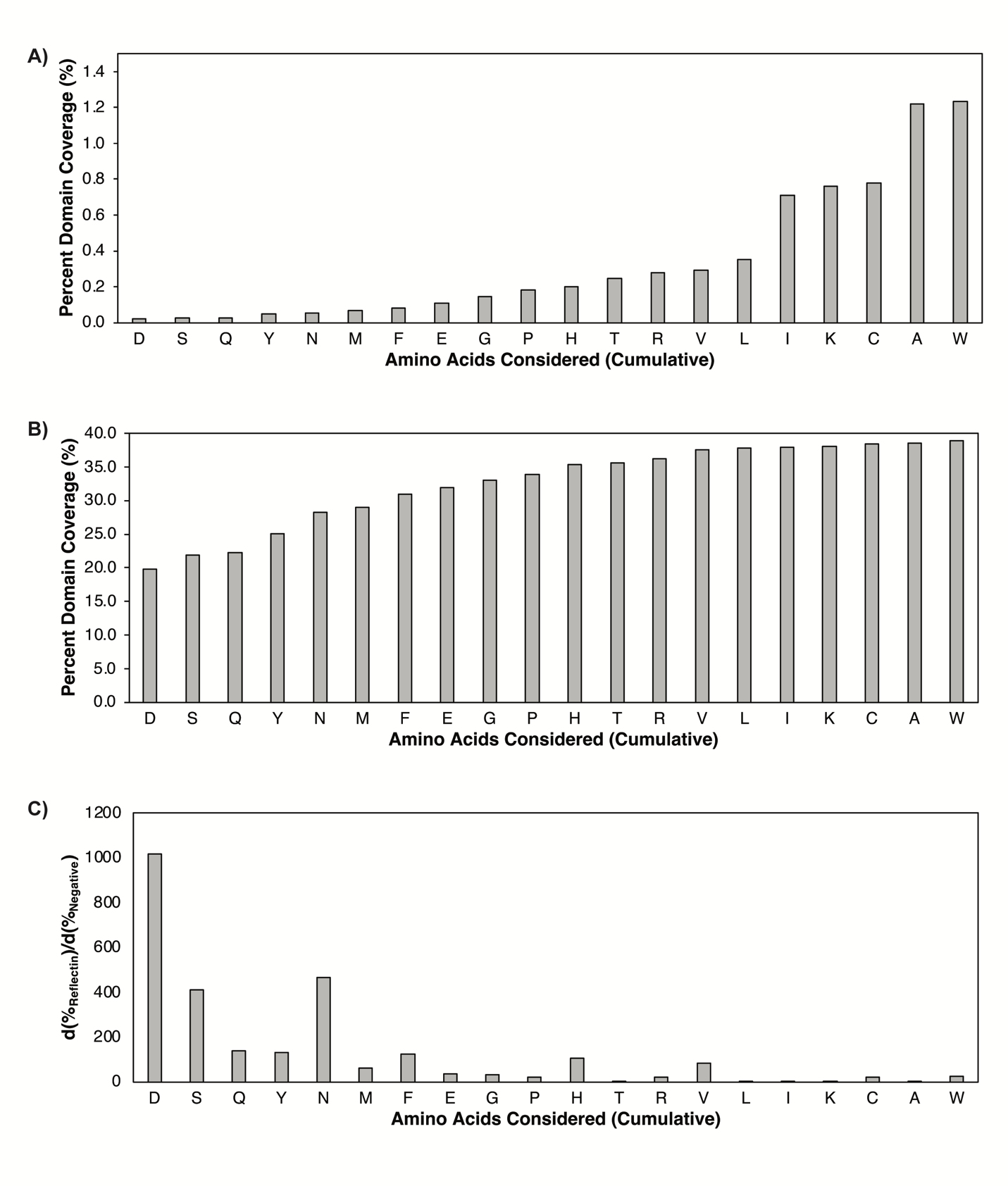

**Figure S4.** Determining Z position to eliminate false positives. The negative control set A) had 5000 sequences randomized across cephalopod species, whereas the B) reflectin set had the 141 reflectin sequences. The C) ratio of domain coverage of reflectins and the negative control was used to distinguish which residues can be considered for Z.

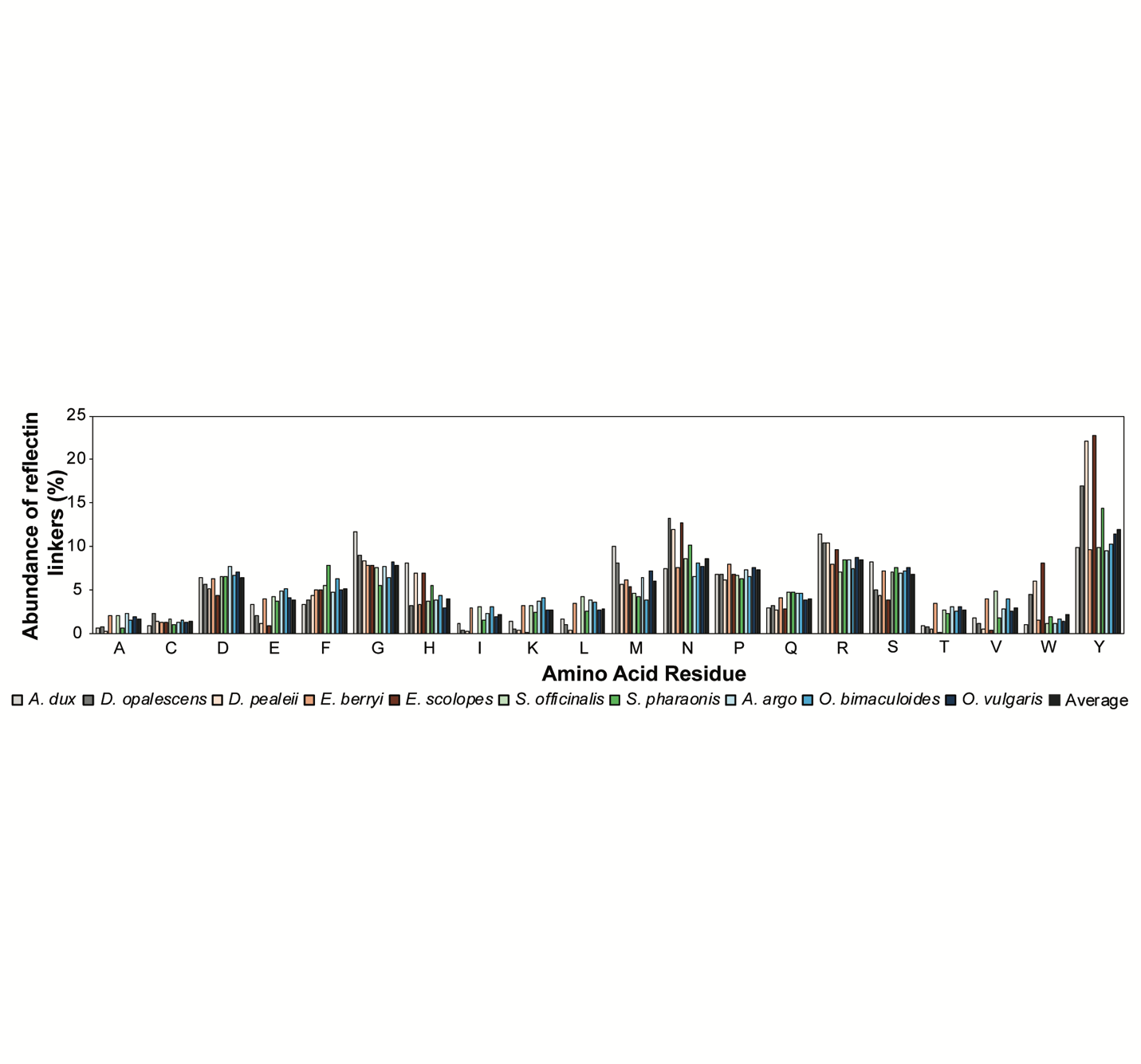

**Figure S5.** Amino acid composition of cephalopod reflectin sequences using our updated model. All sequences were submitted to Protein Calculator v3.4 (protcalc.sourceforge.net) to obtain the counts of amino acid abundance within linkers.
